# The cardiac myosin binding protein-C phosphorylation state as a function of multiple protein kinase and phosphatase activities

**DOI:** 10.1101/2023.02.24.529959

**Authors:** Thomas Kampourakis, Saraswathi Ponnam, Daniel Koch

## Abstract

Phosphorylation of cardiac myosin binding protein-C (cMyBP-C) is a crucial determinant of cardiac myofilament function. Although cMyBP-C phosphorylation by various protein kinases has been extensively studied, the influence of protein phosphatases on cMyBP-C’s multiple phosphorylation sites has remained largely obscure. Here we provide a detailed biochemical characterization of cMyBP-C dephosphorylation by protein phosphatases 1 and 2A (PP1 and PP2A) and develop an integrated kinetic model for cMyBP-C phosphorylation using data for both PP1, PP2A and protein kinases A (PKA), C and RSK2. We find strong site-specificity and a hierarchical mechanism for both phosphatases, proceeding in the opposite direction of sequential phosphorylation by PKA. The model is consistent with published data from human patients and predicts complex non-linear cMyBP-C phosphorylation patterns that are validated experimentally. Our results emphasize the importance of phosphatases for cMyBP-C regulation and prompt us to propose reciprocal relationships between cMyBP-C m-motif conformation, phosphorylation state and myofilament function.

## Introduction

At the molecular level, cardiac muscle function is regulated by several coordinated and periodic processes including electrical excitation and repolarization, calcium cycling, and thick and thin filament activation and their interaction dynamics. Under conditions of physiological stress, humoral and neuronal signals can improve cardiac performance by changing key parameters of these processes which leads to increases in heart rate (chronotropy), force generation (inotropy) and relaxation (lusitropy). Pathological conditions can change these processes, leading to heart disease and heart failure. Often, however, it is not clear whether associated molecular changes are cause or effect of the dysfunction at the cellular or organ level (or, in fact, both).

As a component of the sarcomere, the cardiac isoform of myosin binding protein-C (cMyBP-C) is involved in regulating heart muscle contractility via binding of its N-terminal domains to the actin-containing thin and myosin-containing thick filaments, controlling their regulatory states (1-5). Thin filament binding has been generally associated with an activating effect on contractility by increasing its calcium sensitivity, whereas thick filament binding has been shown to stabilize the myosin heads OFF state and reduce contractility. Physiologically the interaction of cMyBP-C with both filament systems is controlled by its phosphorylation level on several sites in the cardiac-specific m-motif by physiological and pathophysiologically relevant protein kinases (6, 7).

For example, phosphorylation of cMyBP-C by protein kinase A (PKA) leads to enhanced inotropy and lusitropy, making cMyBP-C an important mediator of β-adrenergic stimulation on the level of the contractile myofilaments (8-10). The relevance of cMyBP-C for normal performance and energy efficiency is further emphasized by studies showing that more than 50% of patients suffering from familial Hypertrophic Cardiomyopathy (HCM) carry mutations in the gene encoding for cMyBP-C (11). In addition, cMyBP-C phosphorylation is found to be strongly reduced during heart failure (HF), potentially being both a cause and effect of cardiac dysfunction (12). In good agreement, ablation of cMyBP-C phosphorylation leads to cardiomyopathy and heart failure in transgenic animal models, further underlining its functional significance (13).

A major obstacle to understanding the role of cMyBP-C is posed by its multiple phosphorylation sites and the functional discrepancy between cMyBP-C phosphorylation and phosphomimetic models, making the interpretation of the phenotype of genetic rodent model with e.g. serine-to-aspartate substitutions difficult (14-16). This situation is worsened by the multiplicity of enzymes involved in regulating these sites, including protein kinases A (PKA), protein kinase C (PKC), protein kinase D (PKD), ribosomal S6 kinase (RSK2), Ca^2+^/calmodulin-dependent protein kinase II (CaMKII) and glycogen-synthetase kinase 3 (GSK3) (**Figure 1A**) (reviewed in (7)).

**Figure 1.**
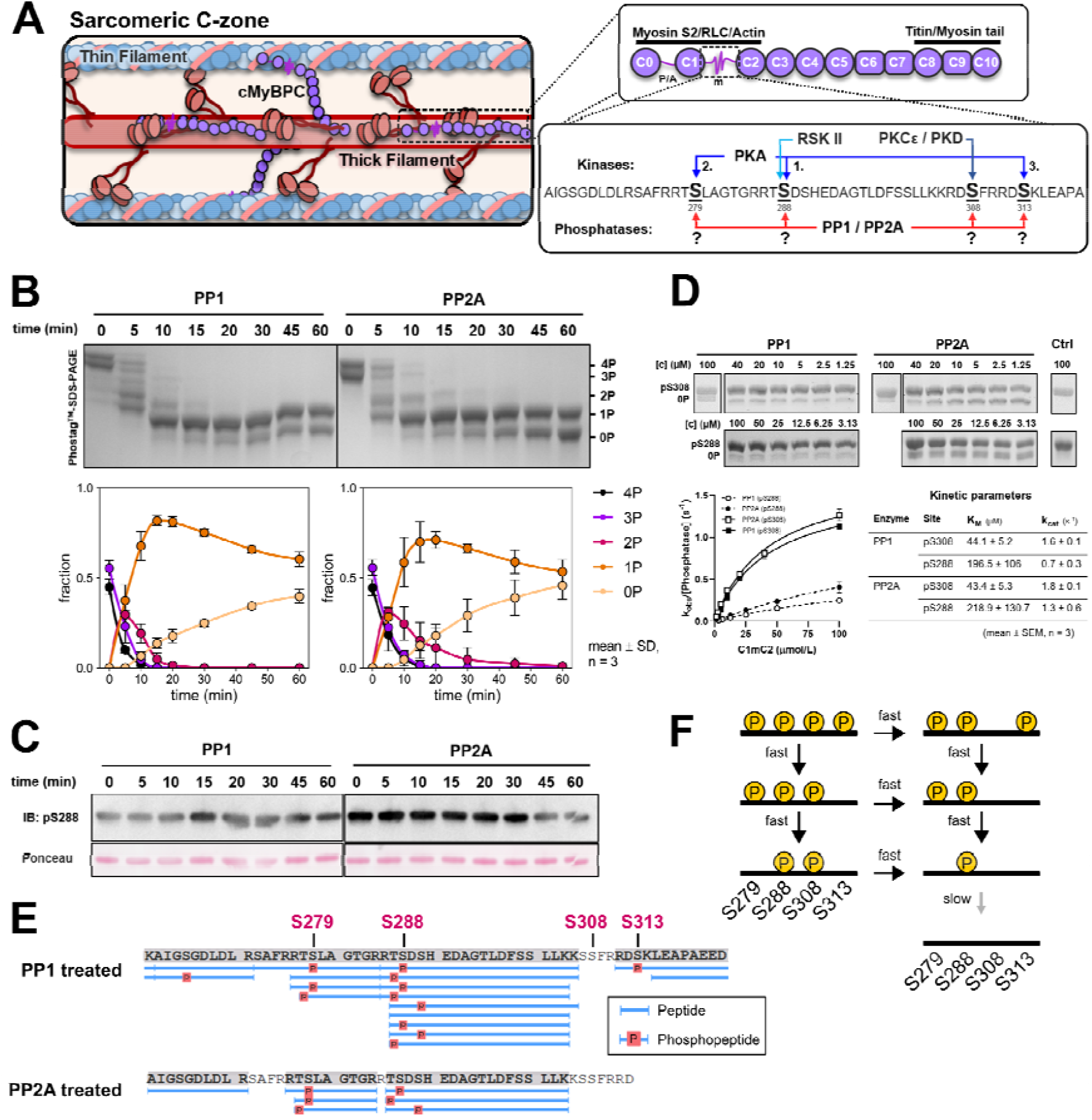
cMyBP-C dephosphorylation by both PP1 and PP2A. **A**, *left and top right:* cMyBP-C is located in the c-zone of the sarcomere. It is composed of eight Ig-, three FnIII-domains, a P/A linker region between C0 and C1 and the semi-structured m-motif between domains C1 and C2 which features multiple phosphorylation sites. The C-terminus is integrated into the thick filament, whereas the N-terminus can bind both to the thin filament, stabilizing the ON-state, and to the thick filament, stabilizing its OFF-state. The phosphorylation states of the m-motif between cMyBP-C C1 and C2 regulate these interactions in a site-specific manner. *Bottom right*: cMyBP-C can be phosphorylated by multiple kinases including PKA, CaMKII (not shown), PKCε and PKD in a site-specific manner. Furthermore, some of these sites influence the phosphorylation state of others. While both PP1 and PP2A have been reported to dephosphorylate cMyBP-C, the reaction mechanism, the site specificity and the kinetics of these reactions have not been characterized yet. **B**, 20μM 3P/4P-C1mC2 (∼50% pS279, pS288, pS308 and ∼50% pS279, pS288, pS308, pS313) were dephoshorylated by 100 nmol/L PP1 or PP2A for 60 min. Samples were taken at different time points and analysed by PhosTag™ -SDS-PAGE. **C**, Immunoblot analysis of samples from B using a pS288-cMyBP-C specific antibody. **D**, Michaelis-Menten kinetics for dephosphorylation of pS288- and pS308-C1mC2 by PP1 and PP2A. **E**, Phosphopeptide specific mass spectrometry analysis of 2P-bands from PhosTag™ SDS-PAGE gels. Clustering of phosphopeptides with multiple apparently different phosphor-residues adjacent to S279 and S288 is likely an artefact resulting from limited resolution. Phosphopeptides from each cluster likely correspond either to pS279 or pS288. **F**, proposed reaction scheme for dephosphorylation of cMyBP-C by PP1 and PP2A. pS279, pS288 and pS313 sites are dephosphorylated sequentially in the reverse order in which they are added by PKA, whereas pS308 can independently be dephosphorylated at any point.

Generally, the phosphorylation state of any protein is a function of both kinases and phosphatases with both enzyme classes being equally important. In the case of multisite and multi-enzyme phosphorylation systems, the relationship between enzyme activity and phosphorylation states is often non-linear and can give rise to complex behaviour such as oscillations or multistability which require quantitative approaches to understand the system behavior (17). Understanding these relationships quantitatively, however, is necessary to pinpoint the physiological roles of a protein’s phosphorylation states. While significant progress has been made in determining the identity and site-specificity of the kinases regulating cMyBP-C (6, 7), very little is known of how cMyBP-C is dephosphorylated. Previous studies have shown that cMyBP-C can be dephosphorylated by both protein phosphatase 1 (PP1) and protein phosphatase 2A (PP2A) *in vitro* (18) and *in vivo* (19, 20), but it is unknown what the site-specificity and kinetics of these dephosphorylation reactions are.

Here we provide a detailed biochemical characterization of the dephosphorylation of cMyBP-C by both PP1 and PP2A, and develop a kinetic model for cMyBP-C phosphorylation that integrates the activities of both kinases (PKA, RSK2, PKC) and phosphatases (PP1 and PP2A). Our model is calibrated using data from dozens of kinetic experiments and further validated by independent biochemical experiments. Our findings show that PP1 and PP2A follow a strongly sequential mechanism for cMyBP-C dephosphorylation that likely depends on intrinsic structural transitions in the m-motif. Individual sites and phosphatases show different dephosphorylation kinetics leading to unique and non-linear steady-state distributions of cMyBP-C phosphorylation states with a Hill-coefficient of 2 for higher phosphorylated forms. We show that our model is consistent with data on cMyBP-C phosphorylation from hearts of healthy donors and HF patients but that the observed changes likely require more complex remodeling in addition to β-adrenergic receptor desensitization and increases in total phosphatase activity during HF (e.g. changes in the PP1/PP2A ratio). Our findings will help to better disentangle the role of cMyBP-C phosphorylation in health and disease, and may have important implications for the modulation of cMyBP-C phosphorylation during HF.

## Results

### PP1 and PP2A dephosphorylate cMyBP-C in reverse order of PKA phosphorylation

To elucidate how and at which sites PP1 and PP2A can dephosphorylate cMyBP-C, we dephosphorylated an almost completely phosphorylated fragment of cMyBP-C containing the m-motif flanked by domains C1 and C2 (C1mC2) (mixture of ∼50% pS279, pS288, pS308 and ∼50% pS279, pS288, pS308, pS313; *rattus norvegicus* sequence) with both protein phosphatases. We found that after an initial phase of rapid dephosphorylation of the tetrakis-, tris- and bis-phosphorylated species by both PP1 and PP2A, dephosphorylation slows down in the presence of only mono-phosphorylated C1mC2 (**Figure 1B**). However, the accumulation of completely dephosphorylated C1mC2 indicates that both PP1 and PP2A can dephosphorylate C1mC2 at all sites, although at least one site appears to be dephosphorylated significantly more slowly than the others.

We have previously shown that PKA phosphorylates C1mC2 in a strictly sequential fashion (pS288 ➔ pS279 ➔ pS313), whereas ribosomal S6 kinase II (RSK2) and PKCε/PKD phosphorylate cMyBP-C site-specifically at S288 and S308, respectively (6). We thus set out to explore the mechanism and kinetics for the dephosphorylation of the individual sites. First, we utilized a pS288-specific antibody and analysed the samples from the dephosphorylation time courses described above. We found that the level of pS288 is relatively constant and well correlated with the mono-phosphorylated species, indicating that dephosphorylation of pS288 likely accounts for the slow dephosphorylation phase (**Figure 1C**). We used Michaelis-Menten kinetics to characterize dephosphorylation of the mono-phosphorylated substrates pS308- and pS288-C1mC2. In agreement with the Western-Blot data, we found that pS288 is a poor substrate for both PP1 and PP2A, exhibiting ∼four-fold higher K_m_ and lower k_cat_ values compared to pS308 (**Figure 1D**). Finally, we determined the identity of the phosphorylation sites in the bis-phosphorylated species of the dephosphorylation time courses using mass spectrometry. We found that for both PP1 and PP2A only pS279 and pS288 can be reliably identified (**Figure 1E**), suggesting that pS308 and pS313 are dephosphorylated before pS279 and pS288.

Taken together, we propose the following reaction mechanism for cMyBP-C dephosphorylation for both PP1 and PP2A (**Figure 1F**): As pS313 is dephosphorylated before pS279 and pS288, and since pS288 is dephosphorylated very slowly, dephosphorylation of the PKA sites in the absence of pS308 likely occurs predominantly by following the sequence pS313 ➔ pS279 ➔ pS288, i.e. in the reverse sequence of PKA phosphorylation. While it is possible that some pS288 sites may be removed before pS279, the slow kinetics of pS288 dephosphorylation will make this path less dominant. pS308 is dephosphorylated before pS279 and pS288 in the 3P/4P-C1mC2 dephosphorylation experiments but can also be dephosphorylated efficiently in the absence of other sites. We conclude that pS308 can likely be removed by PP1/PP2A at any point, which is biologically plausible for a site that, independently from PKA, can be regulated by multiple other kinases of the PKC/PKD family (21, 22).

### An integrated quantitative model of cMyBP-C phosphorylation state regulation

Next, we developed an integrated mathematical model of cMyBP-C phosphorylation considering both protein kinase and phosphatase activities. Such an approach is particularly important for multi-site phosphorylation systems such as cMyBP-C, as quantitative differences and crosstalk between sites or enzymes often lead to the emergence of non-linear behaviours such as ultrasensitivity or bistability, with important consequences for cellular information processing (17).

For simplicity reasons we introduce a new nomenclature for cMyBP-C phosphorylation states that makes such references more convenient and is independent of amino acid numbering across species (**Figure 2A**): we denote rat pS288, pS279 and pS313 by α,β and γ, reflecting the sequential order of their addition by PKA (6), and the PKCε/PKD site pS308 as δ.

**Figure 2.**
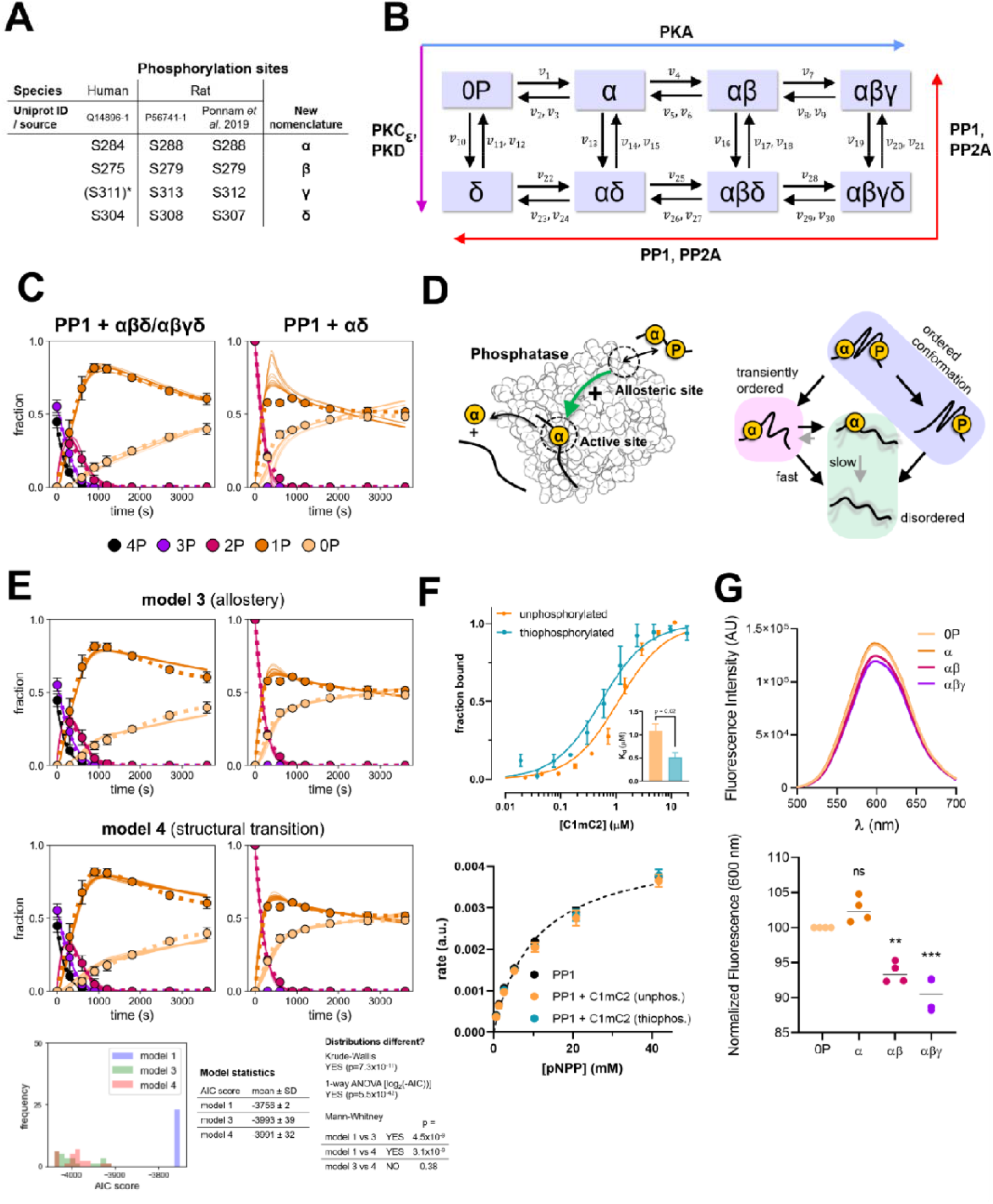
**A**, new nomenclature for cMyBP-C-phosphorylation sites. *Phosphorylation in humans not confirmed. **B**, reaction scheme of proposed cMyBP-C-phosphorylation state model that integrates the activities of multiple kinases and phosphatases acting upon cMyBP-C. **C**, experimental dephosphorylation time course data and model fits show good agreement for 3P/4P-cMyBP-C dephosphorylation, but systematic discrepancies for 2P-cMyBP-C dephosphorylation (here: αδ). **D**, hypothetical mechanisms based on allosterical activation of phosphatases or structural transitions in cMyBP-C might explain the abrupt decline in 1P dephosphorylation rate after 2P-cMyBP-C depletion. **E**, top four panels: improved model fits to the same experimental data as in C following implementation of the allostery or structural transition hypotheses into the model. Bottom panel: model statistics of model 1 (only MM-kinetics), 3 and 4 for PP1 data. Both hypotheses significantly improve model performance according to the Akaike information criterion, but neither hypothesis performs better than the other. **F**, top: binding of unphosphorylated and thiophosphorylated C1mC2 domains to PP1 as measured by microscale-thermophoresis. Bottom: pNPP dephosphorylation kinetics of PP1 in absence or presence of unphosphorylated or thiophosphorylated C1mC2 show no effect of C1mC2 on PP1 activity (n=3). **G**, SYPRO-assay. Top: representative experiment for different phospho-forms of C1mC2. Bottom: quantifcation of peak signal at 600nm of n=4 experiments. ns = not significant, ** p < 0.01, *** p < 0.001.

Based on our mechanistic insights on cMyBP-C (de-)phosphorylation provided here and in earlier studies (6), it is likely that cMyBP-C can adopt eight distinct phosphorylation states as a result of different kinase and phosphatase activities (**Figure 2B**). To keep the model simple and limit the number of parameters, reactions were described by Michaelis-Menten kinetics with substrate competition between sites (23). We collected additional dephosphorylation time course data on different phosphorylation states of C1mC2 for model fitting and a better understanding of the interactions between different sites (**Figure S1**). We calibrated our model using experimental data from the current study and the phosphorylation data previously published using a repeated and combined global/local parameter search approach followed by filtering the best-performing parameter sets (cf. Methods for details). While many individual turnover and Michaelis-constants were not identifiable and varied over >2 orders of magnitudes, specificity constants (k_cat_/K_m_) typically exhibited much lower variability (**Figure S2**). To further increase the robustness of the model predictions, all simulations were performed with multiple independent parameter sets.

The developed model reproduces both the PKA phosphorylation and PP1/PP2A dephosphorylation time courses for substrates with ≥3 phosphate groups. However, we found systematic discrepancies between model output and experimental data for dephosphorylation of bis-phosphorylated forms of C1mC2 (**Figures 2C and S3-4**). Upon closer inspection of the experimental time course data for dephosphorylation of both αβ and αδ (**Figures 2C and S3**), it appears as if the accumulation of completely dephosphorylated C1mC2 suddenly halts in the absence of the bis-phosphorylated C1mC2, despite ample supply of mono-phosphorylated substrate for the phosphatases. Because dephosphorylation of mono-phosphorylated C1mC2 appears to directly depend on the presence of bisphosphorylated C1mC2, we hypothesized that 2P-C1mC2 may directly stimulate the dephosphorylation of 1P-C1mC2.

Implementing this assumption into a simple phenomenological model (**model 2**) improved the quality of the fits (**Figures S5-7**). How might such direct dependence of the α-dephosphorylation rate on 2P-C1mC2 work mechanistically?

One potential mechanism is allosteric activation of PP1 and PP2A by phosphorylated (≥2P) C1mC2 (**Figure 2D, model 3**). Allosteric activation of both PP1 and PP2A by peptides or small molecules has been described previously (24, 25), and cMyBP-C features an RVXF-motif close to its phosphorylation sites (**Figure S8**) which might bind to an allosteric site of PP1 (24). Alternatively, the dephosphorylation mechanism might be regulated by the conformational states of the substrate. In agreement, PKA phosphorylation of cMyBP-C’s m-motif has been reported to induce a disorder-to-order transition (26, 27). We therefore speculated that dephosphorylation of either the δ- or β-site in a bis-phosphorylated C1mC2 may transiently keep an ordered conformation that allows for rapid dephosphorylation of the α-site before collapsing into a disordered state in which dephosphorylation of the α-site is strongly inhibited (**Figure 2D, model 4**). Both mechanistic possibilities considerably improved the model fit to the experimental data in comparison to the initial model and score better when applying the Akaike information criterion (AIC) with, however, similar AIC scores (**Figure 2E and S9-10**). We therefore experimentally tested both hypotheses.

Using pulldown-assays and Microscale Thermophoresis, we confirmed that C1mC2 can bind PP1 with a high affinity, which is further increased upon thiophosphorylation (**Figure 2F** top and **S11**). However, pre-incubation of PP1 with either thiophosphorylated or Ser-to-Asp substituted C1mC2 to mimic phosphorylation did not increase PP1 activity in both para-Nitrophenylphosphate (pNPP) assay or towards tetrakis-phosphorylated C1mC2 (**Figure 2F**, bottom and **Figure S12**), suggesting that higher phosphorylated forms (≥2P) of C1mC2 do not allosterically activate PP1.

To experimentally test the structural transition hypothesis (model 4) would require simultaneous measurements of dephosphorylation time-courses and structural conformation of C1mC2. However, this type of experiment would only be feasible with technical developments beyond the scope of the present study. We thus sought to obtain experimental evidence using a steady-state approach. If our hypothesis is correct, phosphorylation of C1mC2 at the α-site alone would not be sufficient to push the conformational equilibrium towards an ordered state. To test this prediction, we used the SYPRO-assay. The change in fluorescence intensity upon binding of this dye to hydrophobic surface patches is used in thermal denaturation assays to measure protein stability (28). Since denaturation represents an extreme example of conformational order to disorder transition, we reasoned that the signal from the SYPRO-assay could be used as a proxy for conformational order. We found that αβ- and αβγ-phosphorylation of C1mC2 led to a significant signal decrease compared to the unphosphorylated protein, suggesting a disorder-to-order transition. In contrast, and in agreement with our prediction, we found that phosphorylation of C1mC2 in the α-site did not result in a significant signal change (**Figure 2G**), indicating that the α-site alone cannot induce a significant conformational change. Taken together, these experiments suggest that the structural transition model is more likely to be correct than the allosteric activation model.

### Dose-response and dynamics of cMyBP-C phosphorylation in presence of kinases and phosphatases

Since the structural transition model outperforms the other hypotheses, we implemented this mechanistic assumption for the dephosphorylation of cMyBP-C by both PP1 and PP2A, and calibrated the full model with all datasets. As expected, the updated model performs significantly better than the original Michaelis-Menten kinetics-based model trained to the complete dataset (**Figures S13-15**). To complete the model, we added reactions for RSK2 and PKCε and determined their reaction parameters towards C1mC2 experimentally using Michaelis-Menten kinetics (**Figures S16-S17**). *In vivo*, the phosphorylation state of any protein is a function of the joint activities of kinases and phosphatases and the cross-talk, competition and feedbacks between enzymes and sites often leads to unexpected phenomena.

First, we examined the steady-state phosphorylation of cMyBPC in the presence of kinases and phosphatases. Increasing concentrations of PKA lead to a gradual decrease of unphosphorylated cMyBP-C (0P), concomitant by a broad and high peak of 1P, followed by a small peak of 2P and a switch-like transition to 3P cMyBP-C (**Figure 3A**). Due to the hierarchical nature of cMyBP-C (de-) phosphorylation by PKA and PP1, these species correspond strictly to α, αβ and αβγ. Apart from an expected shift towards high [PKA], increasing PP1 concentrations seem to exert little influence on the qualitative nature of these responses.

**Figure 3.**
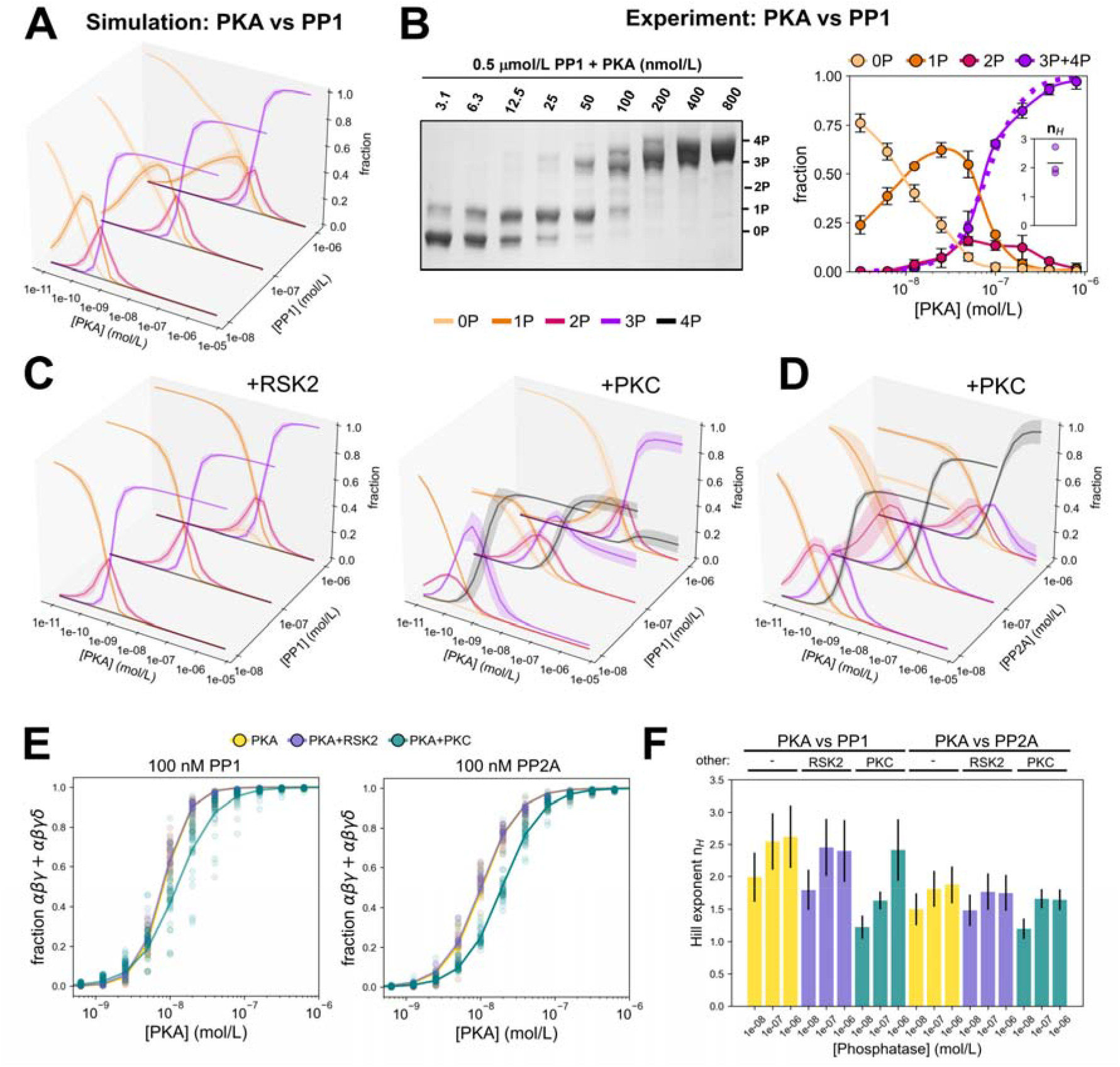
cMyBP-C phosphorylation in presence of kinases and phosphatases. **A**, predicted steady-state cMyBP-C phosphorylation response to increasing PKA concentrations and at different concentrations of PP1 (0P, yellow; 1P, orange; 2P, pink; 3P, purple). **B**, left: experimental steady-state C1mC2 phosphorylation levels in the presence of 0.5 μmol/L PP1 and increasing concentrations of PKA analysed by Phostag-SDS-PAGE. Right: Means ± SD for n=3 repeats. The dashed line indicates the average fit of lumped 3P+4P data using the Hill-equations. Inset: fitted Hill coefficients. **C**, predicted steady-state cMyBP-C phosphorylation levels determined like in A but in the presence of additionally 100 nmol/L RSK2 (left) or PKC (right). **D**, predicted steady-state cMyBP-C phosphorylation levels determined like in A but with PP2A and in the presence of additionally 100 nmol/L PKC (right). **E**, steady-state dose responses of lumped cMyBP-C αβγ+αβγδ states to increasing PKA concentrations at a phosphatase, RSK2 or PKC concentration of 100 nmol/L were fitted to a Hill-equation to quantify the response. Continuous line shows the fitted average. **F**, extracted Hill coefficient parameters of the predicted PKA-dependent steady-state dose-responses of αβγ+αβγδ cMyBP-C for PP1 and PP2A in the presence or absence of PKC or RSK2. Mean±SD of simulations for n=35 parameter sets.

We experimentally tested the model predictions by incubating C1mC2 in the presence of 0.5μmol/L PP1 with increasing concentrations of PKA and analysed the steady-state C1mC2 phosphorylation levels using Phostag™-SDS-PAGE. As shown in **Figure 3B**, the experimental data are qualitatively in very good agreement with the model predictions, showing a gradual decrease in 0P C1mC2 followed by broad and small peaks of 1P and 2P, respectively, and a rapid transition to higher phosphorylated species (3P and 4P). In fact, both the model and experimental data show a switch-like transition from mainly 1P to a mainly 3P species within a narrow range of PKA activity (10-100 nM) that is well described by the Hill equation with a Hill coefficient (n_H_) of ≈ 2 (**Figure 3B, inset** and **3F**).

Next, we examined the steady-state cMyBP-C phosphorylation dose-responses of PKCε and RSK2 in presence of PP1 which showed the expected behaviour of single-site phosphorylation systems (**Figure S18**). Due to the reported cross-talk between sites (6), we also studied the responses to PKA in presence of RSK2 and PKCε (**Figure 3C**). In all simulations, the presence of 100 nM RSK2 is sufficient to convert all cMyBP-C into the 1P (α)-state. Further increasing PKA concentration resulted in a gradual decrease of 1P and small 2P (αβ) peak followed by the complete conversion of cMyBP-C into the αβγ state.

The general qualitative features of all responses considered so far are preserved between PP1 and PP2A (**Figure S18-19**), although the transition to the αβγ state in response to PKA appears to be more switch-like in the case of PP1. The presence of 100 nM PKCε, however, revealed interesting differences between PP1 and PP2A: if PP1 is present, the endpoints of the 4P- and the biphasic 3P-cMyBP-C curves in response to increasing PKA concentrations strongly depend on the total PP1 concentration (**Figure 3C**, right). In contrast, increasing PKA in the presence of PP2A leads to transient intermediate 2P and 3P peaks followed by the near-complete transition to the 4P state for all PP2A concentrations (**Figure 3D**). While PKCε dependent cMyBP-C phosphorylation at the δ-site impairs PKA-dependent phosphorylation of the γ-site, PKCε dependent phosphorylation of cMyBP-C has been reported to exert only minor effects on thin filament structure and no effects on thick-filament structure and force generation (6). Phosphorylation of the δ-site therefore likely modifies other cMyBP-C phosphorylation sites without directly influencing sarcomere function itself.

Thus, we re-analysed the simulation data by lumping together all phosphorylation states which only differ by the presence or absence of the δ-site phosphate (**Figure S20**). Surprisingly, however, qualitative differences between PP1 and PP2A remained. In particular, at low PP1 concentrations, high levels of αβ+αβδ (>50%) can be reached, whereas at high PP1 concentration, cMyBP-C transitions almost directly from a dominant 0P+δ state to the αβγ+αβγδ state upon increasing PKA (**Figure S20, left**). For PP2A, the system transitions invariably from a moderate 0P+δ state through a small region where αβ+αβδ dominates to αβγ+αβγδ (**Figure S20**, right).

Since the transition to the αβγ and/or αβγδ state in response to PKA often appeared more switch-like for PP1 compared to PP2A, we quantified these properties by fitting the αβγ+αβγδ responses to a Hill-equation to extract Hill-exponents that describe the steepness of the response. Interestingly, at 100 nmol/L phosphatases, addition of PKCε seemed to decrease the steepness of the PKA responses in presence of PP1 (**Figure 3E, left**), but only led to a rightwards shift of the curve in presence of PP2A without affecting apparent cooperativity (**Figure 3E, right)**. Indeed, we found that Hill-coefficients of the αβγ+αβγδ response for PP1 were generally larger (ranging from ∼1.25 – >2.5) than for PP2A (ranging from ∼1.25 – <2) (**Figure 3F**). In contrast, EC_50_ values of the same responses tended to be higher for PP2A than for PP1. Interestingly, PKCε phosphorylation mainly shifts the EC_50_ of the αβγ+αβγδ response when PP2A is present, whereas for PP1, PKC phosphorylation leads to a Hill-coefficient that monotonically increases with PP1 concentration (**Figure 3F**). A full summary of the statistical analyses of these quantities is given in **Table S1**. We conclude that a major function of the δ-site is likely to modify the responsiveness of the remaining sites to PKA phosphorylation.

We also examined the dynamic behaviour of the model in response to changes in PKA activity in presence of both PP1 and PP2A (please see **Supplemental Information Text** and **Figure S21**). Of note, dephosphorylation time-courses following kinase removal converged on a slowly decreasing 1P trajectory that only depends on the total phosphatase concentration and likely results from the slow α-cMyBP-C dephosphorylation kinetics.

### Potential effects of heart failure associated changes in PKA and phosphatase activities on cMyBP-C phosphorylation

cMyBP-C phosphorylation has been reported to be significantly reduced during HF, likely reflecting β-adrenergic receptor desensitization and increases in total phosphatase activity (12, 29-32). To see whether our model, whose kinetic parameters were calibrated on *in vitro* data only, is consistent with the cMyBP-C phosphorylation state distribution data from non-failing or failing donor hearts reported previously (12), we initially fitted our model to the non-failing data set using only the five enzyme concentrations of PKA, PKCε, RSK2, PP1 and PP2A (restricted to a physiologically plausible range), but not the kinetic constants as free parameters. The fitted distributions largely capture the relationships between cMyBP-C phosphorylation states that are experimentally observed in non-failing donor hearts (**Figure 4A** left, donor (CL) vs donor (fit)), suggesting that these concentrations might correspond to the baseline enzyme activities observed in non-failing hearts.

**Figure 4.**
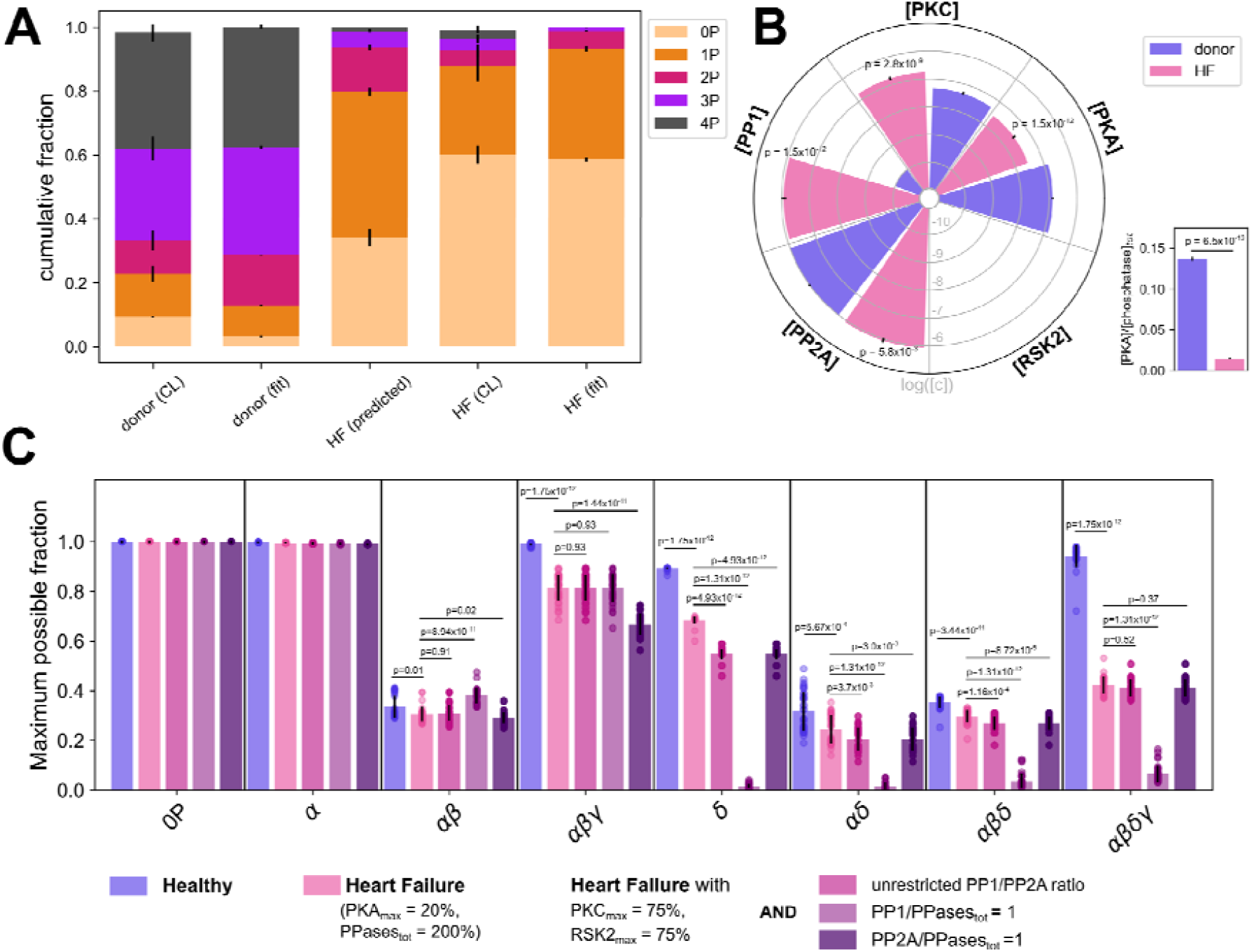
cMyBP-C phosphorylation states during heart failure (HF). **A**, the model was tested for consistency with experimental data on cMyBP-C basal phosphorylation states in hearts from healthy donors or HF patients reported in Copeland et al. 2010 (CL) by fitting the model using only the enzyme concentrations as free parameters (fit). Additionally, the distribution of cMyBP-C phosphorylation states during HF was predicted by starting with the enzyme concentrations fitted to the donor (CL) data followed by a decrease in [PKA] by 50% and a 2-fold increase of total phosphatase concentrations consistent with previous reports on β-adrenergic receptor downregulation and phosphatase activity during HF. **B**, comparison of the fitted enzyme concentrations underlying donor (fit) and HF (fit) data from **A** and calculated PKA/phosphatase_tot_ ratios. **C**, Maximally achievable fraction for each cMyBP-C phosphorylation state under various conditions. For each phosphorylation state, algorithmic optimization was used to find the enzyme vector [PKA, PKC, RSK2, PP1, PP2A] (within a physiological range) that maximizes the phosphorylation state under consideration. To probe the effect of perturbing other enzymes during HF, PKC and RSK2 concentrations were further restricted and PP1/PP2A ratio was either allowed to vary freely (crimson), or phosphatases were fixed to PP2A/PPasetot = 1 (purple) or PP1/PPasetot = 1 (dark purple). Each fitting or optimization run has been performed with all of the n=35 parameter set. Values represent mean ± SEM.

During HF, β-adrenergic receptors are strongly desensitized and total phosphatase activity in cardiomyocytes increases roughly two-fold during HF (30, 31). To determine whether these changes are sufficient to explain the observed changes in cMyBP-C phosphorylation during HF, we tried to predict cMyBP-C phosphorylation during HF as follows: we used the enzyme concentrations obtained by fitting the model to the data from non-failing heart donors, decreased PKA concentration by 50% and increased total phosphatase concentration by two-fold (i.e. constituting a 4-fold decrease in PKA/phosphatase ratio). Interestingly, the predicted pattern is at odds with the experimental one and overestimates the amount of cMyBP-C phosphorylation (**Figure 4A**, HF (predicted) vs HF (CL)). However, since it is difficult to estimate the decrease in PKA activity during HF from receptor desensitization alone and changes in other kinase activities might contribute to HF-related changes in cMyBP-C phosphorylation, we also fitted the model directly to the failing heart data while keeping the two-fold increase in total phosphatase activity as a constraint. As shown in **Figure 4A** (HF (CL) vs HF (fit)), the fitted model shows good agreement with the data from failing heart samples. Interestingly, when comparing the estimated enzyme concentrations for the non-failing and failing donor hearts, we indeed found further differences in addition to the expected decrease in PKA such as a significant increase in PKCε concentration by a factor of ∼4 (**Figure 4B**). Most strikingly, the two-fold increase in total phosphatase concentration was almost exclusively a result of increased PP1, whereas PP2A barely changed. For non-failing donor hearts, in contrast, the fitted PP1 concentration was very low compared to the PP2A concentration. To gain further insights into the relevance of these predictions, we performed a control experiment by fixing the PP1/PP2A ratio to PP1/PPases_tot_=1 for donor hearts and to PP2A/PPases_tot_=1 for failing hearts during the fitting procedure. Interestingly, we found the quality of the fits to be notably reduced (**Figure S22**), indicating that the ratio between these phosphatases is an important determinant of the cMyBP-C phosphorylation state. On a side node, the fitted enzyme concentrations amount to a 10x decreased PKA/phosphatase ratio (**Figure 4B**, right), thus explaining why a 4x reduction in PKA/phosphatase ratio was insufficient to predict cMyBP-C patterns in failing donor hearts. Lastly, the fitted RSK2 concentrations shown in **Figure 4B** remained at zero in both conditions, indicating that RSK2 is not required to explain any of the human heart data.

These findings indicate that the remodeling process of the signaling networks controlling cMyBP-C phosphorylation during HF could be more complex than previously assumed. Specifically, we wondered whether changes in PKCε activity or in the ratio of PP1 to PP2A might even have beneficial effects by allowing cMyBP-C to access phosphorylation states that otherwise might be precluded by β-adrenergic receptor desensitization and increases of total phosphatase activity. To address this issue and gain more functional insights, we rephrased this question as an optimization problem and asked: to which degree can a cMyBP-C phosphorylation state of interest be maximised and what are the required enzyme concentrations?

Guided by the concentrations resulting from fitting the model to donor heart data, we identified the maximally achievable fractions (MAFs) for each of the eight cMyBP-C phosphorylation states and their corresponding enzyme vectors under various conditions **(Figure 4C** and **S22-26)**. In agreement with our dose-response data, we found that the MAFs for intermediate states (αβ, αδ, αβδ) are generally lower than for other states. Under HF conditions, the MAFs of most states were unaffected, but moderate to strong reductions were found for αβ, αβγ and αβγδ, consistent with the strongly reduced amount of 3P and 4P cMyBP-C in the failing donor hearts (**Figure 4C**, blue vs red). To better understand the contribution of individual enzymes that might be altered by HF-related remodeling, we repeated the optimization analysis by further restricting the maximum concentrations of RSK2 and PKCε and/or by clamping the relative amounts of PP1 and PP2A (without changing total phosphatase concentrations). Interestingly, while reducing RSK2 and PKCε showed little effect (except on δ), a significant reduction of αβγ could be seen when fixing PP2A/PPase_tot_ to 1, and almost complete abolishment of δ, αδ, αβδ and αβγδ when fixing PP1/PPase_tot_ to 1. These findings highlight the functional importance of phosphatases for regulation of individual cMyBP-C phosphorylation states and show that changes in the PP1/PP2A ratio may indeed compensate some of the HF-associated changes irrespective of total phosphatase concentration.

## Discussion

The results presented above show that the phosphorylation state of cMyBP-C, an important regulator of contractile function in both the healthy and diseased heart, is strongly dependent on *both* protein kinase and phosphatase activities. Although the former has been extensively studied, the activity, selectivity and regulation of individual phosphatases targeting cMyBP-C have remained largely elusive or contradictory (18-20, 33). In the present work, we show that both PP1 and PP2A dephosphorylate cMyBP-C on several key sites in a highly regulated manner *in vitro*, likely controlled by intrinsic structural changes in cMyBP-C phosphorylation motif itself (26, 27). We also report a novel interaction between PP1 and cMyBP-C that does not depend on cMyBP-C being phosphorylated but likely on its RVxF motif. Strikingly, the dephosphorylation of m-motif serines by both PP1 and PP2A obeys a hierarchical mechanism following the reverse order of their phosphorylation by PKA (6, 34).

The kinetics and the hierarchical phosphorylation/dephosphorylation mechanism have important implications for dynamic changes in cMyBP-C phosphorylation and associated sarcomere function. Rapid phosphorylation of the initial α-site by PKA coupled to slow dephosphorylation kinetics by phosphatases (after removal of the higher phosphorylation states) might constitute a form of biochemical memory. In agreement with this, our model predicts significant phosphorylation of the α-site after removal of PKA activity for several minutes to hours (Figure S21), depending on the phosphatase concentration. Moreover, since the α-site phosphorylation is a pre-requisite for phosphorylation of any subsequent site, a biochemical memory encoded in the α-site phosphorylation might potentiate cMyBP-C phosphorylation during repeated, submaximal β-adrenergic stimulation. In fact, a similar mechanism has been proposed for the post-tetanic potentiation of skeletal muscle via phosphorylation of the myosin regulatory light chain (35).

Based on our previously published phosphorylation data (6) and the results of our current biochemical studies using both isolated PP1 and PP2A, we have developed a kinetic model of the local (in terms of network topology) cMyBP-C phosphorylation regulatory network integrating both protein kinase and phosphatase activities (Figure 2). Importantly, our model suggests that the phosphorylation/dephosphorylation kinetics of cMyBP-C are controlled via a conformational switch in the phosphorylatable m-motif itself. Although further experimental confirmation is required, the potential reciprocal relationships between m-motif structure, phosphorylation state and sarcomere function raise the intriguing possibility of a mechano-biochemical feedback loop: assuming that the forces developed during the contractile cycle acting on cMyBP-C have the capability of changing the conformation of the m-motif (36), such forces might alter the phosphorylation state of cMyBP-C which, as a key parameter of sarcomere function, would in turn affect force dynamics. In line with this idea, phosphorylation-dependent structural unfolding of the m-motif of cMyBP-C (26, 27) and changes in cMyBP-C phosphorylation following mechanical stress have been reported previously (37, 38). However, the relationship between conformation and phosphorylation seems reciprocal and how exactly force affects m-motif structure and phosphorylation state of cMyBP-C in the intact sarcomere is not yet known, thus the effects and details of such feedback regulation remain to be elucidated. A promising first approach to do so might be to interface our current model with existing models of sarcomere dynamics (36, 39, 40).

We have previously shown that the distinct phosphorylation states of cMyBP-C results in different affinities of its N-terminal domains for thick and thin filaments, coupled to different effects on their respective regulatory states (6, 41). This suggests that cMyBP-C might act as a ‘gear-box’ of the sarcomere, where different combinations of kinase and phosphatase activities lead to distinct phosphorylation states that modulate thin and thick filament function, and optimize their working under different physiological conditions, perhaps ensuring energetic efficiency of the contractions. Indeed, a role for cMyBP-C in contractile energy efficiency is widely discussed (42-44) and the hypothesized mechano-biochemical feedback may be crucial for such a function.

Another important finding of both of our simulations and experiments is the hypersensitivity or apparent cooperativity of the cMyBP-C phosphorylation state (Figure 3), suggesting that the system is exquisitely sensitive to small changes in the PKA/phosphatase activity. As shown in Figure 3A, reducing active PKA concentration by a factor of two leads to a redistribution of the cMyBP-C phosphorylation state from a mostly tetrakis/tris-phosphorylated species (∼70% of total) to an almost entirely mono- and unphosphorylated species. This is further emphasized by the steepness of the [PKA]-[3P/4P] relation with Hill-exponents bigger than two. Since the Hill-exponent does not increase with lower phosphatase concentration (Figure 3F), the effect is likely an intrinsic property of the sequential cMyBP-C (de-)phosphorylation mechanism rather than zero-order ultrasensitivity (45, 46). This suggests that at some threshold sympathetic tone, β-adrenergic signalling likely leads to switch-like transition into a regime in which the majority of C-zone myosin heads become available for cross-bridge cycling. Perhaps such a switch-like transition is one of the main functions of cMyBP-C multisite phosphorylation/dephosphorylation. Similar switch-like responses stemming from multisite (de-)phosphorylation have been described by us for phospholamban (47). In the case of cMyBP-C, however, not all phosphorylation site combinations might correspond to discrete functional states. Some phosphorylation site combinations may be functionally redundant and their transitions more gradual. While further work is necessary to better separate the functional roles of these combinations, our model may be useful to design experiments allowing such functional separation without the need for Ser to Asp/Ala substitutions.

We also examined the validity of our model for cMyBP-C phosphorylation in basal and disease conditions using experimental data from human donors and heart failure patients (12) and found the model to be consistent. Interestingly, although increasing cMyBP-C phosphorylation appears to be a primary mechanism by which the heart activates its contractile reserve under conditions of increased cardiac stress at the level of the sarcomere (48), the basal level of cMyBP-C phosphorylation has been reported to be rather high at about 2-3 mol P_i_/mol MyBP-C (12, 15, 29). Assuming such high basal phosphorylation is correct, this raises two questions: firstly, what is the physiological role and secondly, what is the determinant of high cMyBP-C phosphorylation level under basal conditions? While high basal phosphorylation perhaps allows for a larger range of myofilament regulation, e.g. during parasympathic stimulation (49), our findings suggest that the low substrate specificity of PP1 and PP2A for the α-site may partly contribute to the high basal phosphorylation of cMyBP-C.

Furthermore, our analyses suggest that the observed cMyBP-C hypo-phosphorylation during heart failure might not be solely explained by decreasing PKA and increasing phosphatase activity. Indeed, the biggest contributor to the observed hypo-phosphorylation in our simulations is a shift in the ratio of PP1 to PP2A activity, and changing this ratio can potentially increase cMyBP-C phosphorylation in a heart failure setting. Additionally, our model also predicted PKCε activity acting on cMyBP-C to be significantly elevated. Although the precise role of α-adrenergic signalling for the heart muscle function has remained elusive, both *in vivo* studies in animal models and isolated intact muscle preparations showed a robust positive inotropic response after α1-adrenergic receptor stimulation (50, 51). Our previous study (6) and our current findings highlight a novel crosstalk between α- and β-adrenergic stimulation on the level of the contractile myofilaments. PKCε phosphorylation of cMyBP-C not only increases the EC_50_ for phosphorylation of cMyBP-C by PKA but also reduced its sensitivity (Fig. 3). Thus, α-adrenergic signalling via PKCε might allow a more graded transition of cMyBP-C phosphorylation states, which allows access to intermediate phosphorylation states that are not accessible during only PKA/β-adrenergic stimulation (e.g. 2P states). In good agreement, the bis-phosphorylated state of cMyBP-C is underrepresented under both basal conditions in both human and rodent hearts (12), suggesting that this state might be more accessible during disease conditions. Note, however, that also PKD has been reported to phosphorylate the δ-site of cMyBP-C and could thus play a similar role alternatively or in addition (21). However, while our model predicts that activity changes in multiple enzymes are required to explain the observed cMyBP-C phosphorylation pattern during HF, the reported changes in the literature (particularly for phosphatases PP1 and PP2A) are inconsistent, perhaps reflecting differences in experimental models, aetiology or disease progression (52-55).

An important limitation of our study particularly concerning the interpretation of the human heart data (Figures 3A and 3B) is that the current model is incomplete with respect to components further upstream in the cardiac signalling network (or CamKII, PKD and GSK3 for that matter), some of which likely have important implications for cMyBP-C phosphorylation. In contrast to the *in vitro* situation of our study, many enzyme activities at the sarcomere are regulated in a highly dynamic manner due to regulation by various subunits *in vivo*. Substrate dephosphorylation by PP1, for example, is subject to regulation by feedforward loops involving e.g. PKA, PKC and inhibitor-1 (47, 56), whereas sarcomeric PP2A activity is regulated by B56-subunits e.g. via changed localization after β-adrenergic stimulation (20, 33, 57) and via Pak1 (58), which itself lies downstream of multiple receptor tyrosine kinases. A translocation to the myofilaments in response to β-adrenergic stimulation has also been reported for PKCε (59). Understanding how these dynamic signaling processes impact cMyBP-C phosphorylation in a site-specific manner is, however, poorly understood and requires further experimental work.

In conclusion, we have provided a detailed and site-specific biochemical characterization of cMyBP-C dephosphorylation by PP1 and PP2A and have developed the first integrated model of cMyBP-C phosphorylation as a function of multiple kinases and phosphatases. Our results highlight the importance of phosphatases for regulating cMyBP-C phosphorylation and open up several interesting avenues for future research.

## Material and Methods

### Recombinant proteins

The catalytic subunit 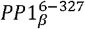 was cloned into a pET15b vector and validated via sequencing by Bio Basic/(USA). For purification of PP1, 50 μL SoluBL21™ Chemically Competent *E. coli* (AMSBIO, UK) cells were transformed with 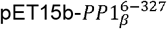 and a 100 mL pre-culture was grown in LB-medium overnight at 37°C in ashaking incubator at 200 rpm. On the following morning 2×1 L autoinduction medium (60) supplemented with 1mM MnCl_2_ were inoculated with 50 mL of the preculture each and further grown at 30°C until the evening before the temperature was lowered to 20°C and cultures were left to grow for ∼72 hrs before harvesting. After harvesting, cells were resuspended in lysis buffer (50 mM TRIS pH 8, 700 mM NaCl, 1mM MnCl2, 1 mM DTT, 25 mM Imidazole, cOmplete™ EDTA-free Protease Inhibitor Cocktail (Roche, cat. no. 11873580001)) and lyzed via 2x freeze/thaw cycles followed by addition of lysozyme, addition of Triton X-100 ad 0.2% and sonication. After clearing the lysate from insoluble material via centrifugation and filtration, PP1 was purified via a HisTrap™ FF column (GE Healthcare, cat. no. 29048609) followed by size exclusion chromatography (Superdex™ 75 pg, GE Healthcare, cat. no. 28-9893-34) into a final buffer of 50 mM TRIS pH 8, 700 mM NaCl, 1mM MnCl_2_, 1 mM DTT. The protein was concentrated to a final concentration of ≈ 0.2 mg/ml (determined by absorbance at 280 nm), aliquoted, snapfrozen in liquid nitrogen and stored at -70°C. To avoid activity loss, PP1 was not refrozen after thawing of aliquots.

The phosphorylated cMyBP-C fragments encompassing domains C1, m-motif and C2 (C1mC2) were prepared as described previously (1, 6).

Protein Phosphatase 2A (PP2A) was purchased from Cayman Chemical (cat. no. 10011237), aliquoted and stored at -70°C until further use. To avoid activity loss, PP2A was not refrozen after thawing of aliquots.

### Pulldown experiments

Pulldown experiments for testing whether the catalytic domain of PP1 and cMyBP-C C1mC2 interact with each other, we performed as follows: first, 50 μL Ni-NTA resin (Qiagen, cat. no. 30410) was washed three times in assay buffer (PBS pH 7.5, 500 mM NaCl, 0.01% tween-20, 40 mM Imidazole, 0.5 mM MnCl_2,_ 2 mM DTT) in 1.5 ml tubes before 100 μL of 1 μM Alexa647-labelled (Thermo Fischer, cat. no. A37573) BSA (negative control for bait) or C1mC2 were added to the resin. Next, 75 μL of 5 μM His6-PP1 were added to resin with A647-C1mC2 or A647-BSA (negative control). In another negative control, 75 μL 1xPBS were added to resin with A647-C1mC2. Tubes were incubated for 45 min at RT in a tube shaker at 1000 rpm before a sample of the supernatant was taken and mixed with 5x SDS sample buffer as a reference for the unbound fraction (SDS sample buffer:sample = 1:5). The supernatant was removed, the resin was washed three times in 750 μL assay buffer and the resin was resuspended in 50 μL 1x SDS sample buffer and boiled for 5 min at 100°C. For SDS-PAGE analysis, 12.5 μL sample/lane were run on a gel. After running the gel, the A647-signal was recorded and the gel was stained with Coomassie solution to obtain the total protein signal (Protein Ark, cat. no. GEN-QC-STAIN-1L).

### Microscale Thermophoresis

For quantifying the interaction strength between PP1 and cMyBP-C, binding affinities were determined via microscale thermophoresis on a Monolith NT.115 instrument (NanoTemper Technologies, Germany). Briefly, serially diluted cMyBP-C C1mC2 was mixed 1:1 with 100 nM Alexa647-PP1 in MST buffer (50mM TRIS-HCl pH 8, 500mM NaCl, 1mM MnCl_2_, 2mM DTT, 0.05% tween-20, MnCl_2_ and DTT were added just before the experiment) and incubated at RT for 5 min before samples were loaded with premium capillaries (NanoTemper Technologies, cat. no. MO-K025). Data were recorded sequentially at 40%, 90% and 95% MST power and 25% laser power. The best data quality was obtained at 90% MST power which was thus used for quantifying the binding affinities using the MO.Affinity Analysis software (NanoTemper Technologies, Germany).

### pNPP assay

The p-nitrophenyl phosphate (pNPP) assay was used to test for potential allosteric activation of PP1 by cMyBP-C C1mC2. Briefly, 100 nM PP1 were preincubated with no additional factors or with 2 μM unphosphorylated or thiophosphorylated cMyBP-C C1mC2 in assay buffer (50mM TRIS-HCl pH 8, 500mM NaCl, 1mM MnCl_2_, 2mM DTT, 0.05% tween-20, MnCl_2_ and DTT were added just before the experiment) for 5 min at 30°C. The preincubated PP1 sample was then mixed 1:1 with a freshly made solution of pNPP (Sigma Aldrich, cat. no. 20-106) in serially diluted assay buffer to reach a final concentration of 50 nM PP1 and ∼1-100 mM pNPP in 96-well microplates (greiner bio-one, cat. no. 675076). pNPP turnover was recorded by measuring absorbance at 405 nm over time in a CLARIOstar microplate reader (BMG LABTECH, Germany). The initial rates obtained from the recorded traces were fitted to the Michaelis-Menten equation to estimate V_max_ and K_m_ values.

### Dephosphorylation assays

Site-specifically phosphorylated C1mC2 fragments were prepared as described previously (6). Phosphorylated C1mC2 was gel-filtered into assay buffer (in mmol/L: 20 MOPS pH 7, 100 KCl, 1 MgCl_2_, 1 MnCl_2_, 2 DTT, 0.05% (v/v) Tween-20) using NAP5 columns (Life Technologies) according to manufacturer’s instructions and protein concentrations adjusted to 20 μmol/L. Reactions were started by adding 0.1 μmol/L PP1 or PP2A, and aliquots quenched at the indicated time points with 5x SDS-PAGE loading buffer and snap frozen in liquid nitrogen. Samples were denatured at 100°C for 3 mins and loaded onto a 12% (v/v) acrylamide SDS-PAGE gel containing 50 μmol/L Phostag™-reagent and 100 μmol/L MnCl_2_. Gels were run at 160 V for 90 mins and protein bands visualized using Coomassie staining.

### Western-blotting

SDS-PAGE samples were run on 4%-20% (v/v) acrylamide gradient gels (BIORAD) and transferred onto nitrocellulose membranes using a Trans-Blot SD Semi-Dry Electrophoretic Transfer Cell (Bio-Rad). Primary incubation took place overnight at 4 °C (anti-pSer282; 1:2,000, ENZO Life Sciences, AL-215-057-R050). Secondary antibody incubation was done at room temperature for 1h (HRP-conjugated goat anti-rabbit IgG, 1:2000 dilution, Bio-Rad, 403005). Membranes were washed with Tris-buffered saline containing 0.05% (v/v) Tween-20, emersed in ECL reagent (BIORAD) and blots developed on a BIORAD imager.

### SYPRO Fluorescence Assay

SYPRO orange was purchased from Life Technologies as a 10000-fold stock solution in DMSO. 20 μmol/L unphosphorylated and phosphorylated C1mC2 fragments were incubated with 20-fold SYPRO Orange for 10 min at 25°C in black 96-well plates (Greiner), and SYPRO Orange fluorescence measured using appropriate excitation and emission filter settings using a ClarioStar plate reader (BMG Labtech).

### Identification of phosphorylation states in 2P cMyBP-C during dephosphorylation using mass spectrometry

To identify the phosphorylation sites that are still phosphorylated in the bisphosphorylated cMyBP-C C1mC2 fragment during the dephosphorylation transient starting from the almost phosphorylated C1mC2 fragment, the dephosphorylation reaction was stopped 3-5 min after addition of PP1 or PP2A by addition of SDS sample buffer and 5 min boiling at 100°C. Samples were separated by PhosTag SDS-PAGE and 2P bands were cut out and sent to the Metabolomics and Proteomics Laboratory of the Bioscience Technology Facility of the University of York for identification of phosphorylation sites. Details on the experimental procedure and the full dataset can be found in the supplementary data file “Phos_ID_D428”.

### Mathematical Models

All models have been formulated using ordinary differential equations that describe the rates of phosphorylation and dephosphorylation of cMyBP-C-forms. Generally, enzymatic reactions were described by Michaelis-Menten-type rate laws that account for the possibility of substrate competition (23) between different phospho-forms 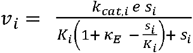, where *e* denotes the concentration of Enzyme E, *s*_*i*_ the concentration ith substrate of E, and *k*_*cat*_, _*i*_ and *K*_*i*_ are the respective turnover and Michaelis-constants and 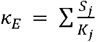 (sum over all substrates *s*_*i*_ of E). For the extended model versions allowing a better description of 2P-cMyBP-C dephosphorylation, the rates were modified as follows:

For the phenomenological model, the rate of α-dephosphorylation was multiplied with (1 + *f*_*act*_ *h*), where the parameter *f*_*act*_ ≥ 0 denotes a maximum activation factor and 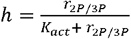 is a hyperbolic function of *r*_*2p/3p*_, the relative amount of bis- and trisphosphorylated cMyBP-C, with a half-saturation constant *K*_*act*_.

For the allosteric activation model, we first derived a general steady-state rate law for allosteric activation for catalysis of a substrate *S*_*j*_ (single site) and multiple non-allosterically competing substrates. Consider the following reaction scheme below where A is an allosteric activator for the catalysis of substrate *S*_*j*_ to product *P*_*j*_ in the presence of multiple competing substrates whose rate of catalysis is not affected by A:

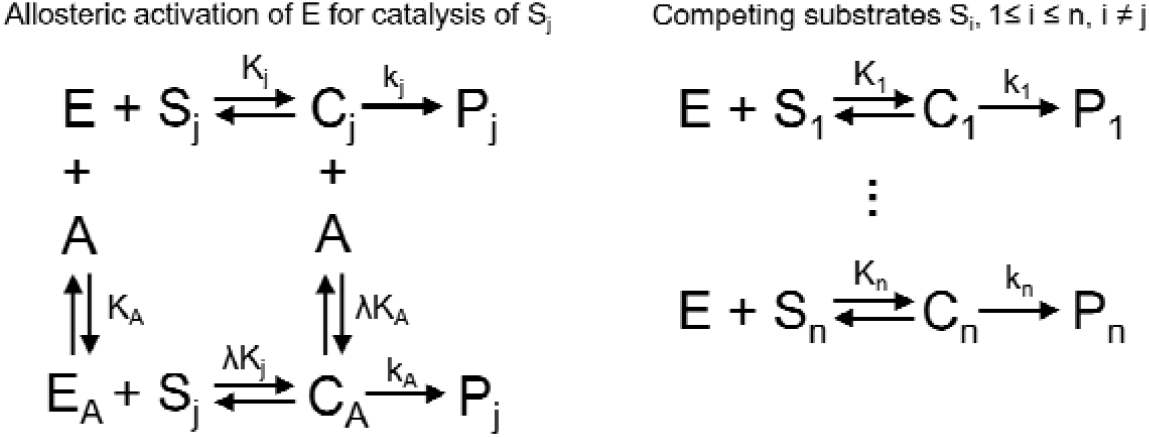

We assume that the total amount of enzyme e_T_ is conserved and given by the relation 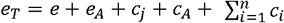. The rate at which product P_j_ is formed is given by *v* = *k*_*j*_ *· e*_*j*_*+k*_*A*_*·e*_*A*_, therefore yielding equation 1: 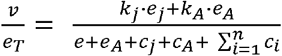. We further assume the rates of complex formation are much faster than catalytic turnover, allowing us to use the rapid equilibrium approximation as outlined in (61) for equilibria K_j_, K_A_, λK_j_, λK_A_ and K_1_, …, K_n_, where λ is a dimensionless scaling parameter to describe the preferential formation of the allosterically activated complex c_A_. Solving the equilibria for the enzyme or complex species and substituting the respective terms in equation 1 yields the final form of the rate law: 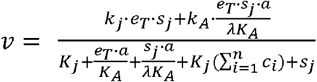. We used this rate law to describe the dephosphorylation of α, assuming for simplicity that only 2P-cMyBP-C species serve as allosteric activators, i.e. A = αβ + αδ.

For the structural transition model, we assumed that dephosphorylation of the β-phosphate from αβ or the δ-phosphate from αδ leads to the production of α’, a (transiently) ordered conformation of α which is a better substrate for phosphatases PP1 and PP2A than α. Isomerization between α’ and α was assumed to follow first order rates V_iso,F_ = k_iso,F_ · α’ and V_iso,R_ = k_iso,R_ · α, respectively.

Further details and ODEs for all models can be found in the Appendix.

For simulation, all models were implemented in Python and integrated with a custom implementation of the fourth-order Runge-Kutta integrator as outlined in (62) with a step-size of 1 second. If necessary, step-size was reduced to 0.1 or 0.01 seconds.

### Model fitting, selection and optimization

For parameter estimation, models were implemented in COPASI v4.35 (63) and parameter estimation was performed with Python using the package basico (v0.4, https://github.com/copasi/basico) (64): For each model, 50-100 independent parameter sets were generated using a global search procedure (Genetic algorithm, generations: 250, population size: 25) and subsequently refined using a local search (Hooke-Jeeves-algorithm, iteration limit 50, tolerance 1e-5, rho 0.2). Generally, weights were assigned by standard deviation and normalized by experiment. However, the following custom weights were assigned manually: PKA-phosphorylation of δ: weight(δ) = 6e-7, weight(αβγδ) = 3e-6. PP1-dephosphorylation of αδ: weight(2P-cMyBP-C) = 6e-7. PP1-dephosphorylation of αβδ/αβγδ and PP2A-dephosphorylation reactions: weight(3P-cMyBP-C) = 1e-5, weight(4P-cMyBP-C) = 1e-5. PP2A-dephosphorylation of αβ and αδ: weight(2P-cMyBP-C) = 5e-6. PP2A-dephosphorylation of αβδ/αβγδ: weight(0P-cMyBP-C) = weight(1P-cMyBP-C) = weight(2P-cMyBP-C) = 2e-6, weight(3P-cMyBP-C) = weight(4P-cMyBP-C) = 1e-5. To filter out parameter sets featuring unrealistic transients resulting from limited temporal resolution in the experimental data, we calculated the sum of squared errors between simulated data and the Akima-interpolation of the experimental data. All parameter sets leading to an error that lies above a predefined cutoff (some cutoffs were set by trial and error, but generally mean + 0.5 standard deviations worked for most conditions) for any of the experiments were filtered out.

For model selection, we utilized the Akaike-Information-Criterion (65). The AIC score for each model and parameter set was calculated as described in (66). A model was judged to be superior to another one if it had a statistically significantly lower AIC score.

For fitting the model to the data from Copeland *et al*. 2010, each of the 35 parameter sets resulting from fitting the model to our *in vitro* data was sequentially applied to a Copasi implementation of the final model which was then used for parameter estimation with enzyme concentrations of PKA, PKC, RSK2, PP1 and total phosphatase concentration as free parameters (Hooke-Jeeves-algorithm, iteration limit 50, tolerance 1e-5, rho 0.2). Based on the previous literature and the kinetic parameters in the present study, enzyme concentrations in the high nanomolar to low micromolar range seemed plausible to us (67). Specifically, parameter search was conducted within the following concentration ranges: PKA ∈ [5e-10, 5e-7], PKC ∈ [0, 2e-7], RSK2 ∈ [0, 2e-7], PP1∈ [1e-10, 1e-6] and total phosphatases ∈ [1e-10, 1e-6] (all in mol/L). PP2A concentration was calculated by total phosphatase concentration minus PP1 concentration, thus the fraction that PP1 or PP2A contributes to total phosphatase concentration was allowed to vary freely. For fitting the heart failure data, the total phosphatase concentration was fixed to twice the total phosphatase concentration obtained by fitting the model to the donor heart data. For control experiments shown in Figure S22, phosphatases were set to either PP1/PPasestot=1 or PP1/PPases_tot_=1.

Similarly, for optimizing individual phosphorylation states, each of the 35 parameter sets resulting from fitting the model to our *in vitro* data was sequentially applied to a Copasi implementation of the final model which was then used for maximizing a chosen cMyBP-C phosphorylation state with enzyme concentrations of PKA, PKC, RSK2, PP1 and total phosphatase concentration as free parameters (Hooke-Jeeves-algorithm, iteration limit 50, tolerance 1e-5, rho 0.2). Permissible enzyme ranges were guided by the resulting concentrations after fitting the model to the data from Copeland *et al*. 2010. More specifically, we assumed that PKA and total phosphatase concentrations resulting from fitting the model to the cMyBP-C phosphorylation data from non-failing hearts (12) represent basal activities in absence of any β-adrenergic or other modulatory stimuli. While we kept total phosphatase concentrations constant, we reasoned that β-adrenergic stimulation might increase PKA concentration by at least 10x or that cholinergic stimulation might further depress basal PKA activity. PKC, RSK2 and PP1/PP2A ratio was subject to the same constraints as before. For optimizing individual cMyBP-C phosphorylation states under conditions resembling a healthy heart, the ranges were thus PKA ∈ [0, 10xPKA_donor(fit)_] ≈ [0, 1.36e-6], PKC ∈ [0, 2e-7], RSK2 ∈ [0, 2e-7], PP1 ∈ [0, PPases_tot,donor(fit)_], total phosphatases = PPases_tot,donor(fit)_ ≈ 1e-6 (all in mol/L). For optimizing individual cMyBP-C phosphorylation states under conditions resembling a failing heart, total phosphatase concentration was increased twofold in line with previous reports (31), while we assumed that the reduction in PKAHF(fit) compared to PKAdonor(fit) at least partly is a result of β-adrenergic receptor desensitization and therefore also applies to conditions of high β-adrenergic stimulation. We thus choose PKA ∈ [0, 10xPKA_HF(fit)_] ≈ [0, 2.89e-7], PKC ∈ [0, 2e-7], RSK2 ∈ [0, 2e-7], PP1 ∈ [0, PPases_tot,HF(fit)_], total phosphatases = PPases_tot,HF(fit)_ ≈ 2e-6 (all in mol/L). To further probe the functional effect of individual enzymes during, we also tested the following conditions (all concentrations in mol/L):

- *<u>HF with PKC/RSK2 restriction:</u>* PKA ∈ [0, 10xPKA_HF(fit)_] ≈ [0, 2.89e-7], PKC ∈ [0, 1.5e-7], RSK2 ∈ [0, 1.5e-7], PP1 ∈ [0, PPases_tot,HF(fit)_], total phosphatases = PPasestot,HF(fit) ≈ 2e-6
- *<u>HF with PKC/RSK2 restriction and clamped PP1:</u>* PKA ∈ [0, 10xPKA_HF(fit)_] ≈ [0, 2.89e-7], PKC ∈ [0, 1.5e-7], RSK2 ∈ [0, 1.5e-7], PP1 = PPases_tot,HF(fit)_, total phosphatases = PPases_tot,HF(fit)_ ≈ 2e-6
- *<u>HF with PKC/RSK2 restriction and clamped PP2A:</u>* PKA ∈ [0, 10xPKA_HF(fit)_] ≈ [0, 2.89e-7], PKC ∈ [0, 1.5e-7], RSK2 ∈ [0, 1.5e-7], PP1 = 0, total phosphatases = PPases_tot,HF(fit)_ ≈ 2e-6
- *<u>No RSK2:</u>* PKA ∈ [0, 10xPKA_donor(fit)_] ≈ [0, 1.36e-6], PKC ∈ [0, 2e-7], RSK2 = 0, PP1 ∈ [0, PPases_tot,donor(fit)_], total phosphatases = PPases_tot,donor(fit)_ ≈ 1e-6

### Statistical analysis

Statistical comparisons between two experimental distributions have been performed in GraphPad prism using a two-sided t-test. Data from simulations were analyzed in Python using the packages numpy (v1.21.5), scipy (stats module, v1.9.3) and statsmodels (v0.13.2). Distributions from simulations often appeared non-normal, which was confirmed using the Shapiro-Wilk test (scipy stats). For comparisons between two non-normally distributed variables, the Mann-Whitney test was used (scipy stats). Comparisons between multiple groups were performed using the Kruskal-Wallis test (scipy stats) and, in case of model selection based on AIC score, additionally by a 1-way ANOVA (scipy stats) on the log_2_-normalized distributions. In case of multiple comparisons, p-values were corrected using the Benjamini/Hochberg procedure with a false discovery rate of 0.05 (statsmodels). For any test, a p-value below 0.05 was considered to indicate a statistically significant difference.

## Code and data availability

All code and quantified raw data used for model calibration will be available upon final publication of the manuscript.

## CRediT author statement

**Thomas Kampourakis**: Conceptualization, Investigation, Writing – Review & Editing. **Saraswathi Ponnam**: Resources, Writing – Review & Editing. **Daniel Koch**: Conceptualization, Supervision, Formal Analysis, Investigation, Writing – Original draft.

## Acknowledgements

We would like to thank Frank Bergmann for help with the basico package. We acknowledge the British Heart foundation and the European Molecular Biology Organization for financial support.

