## Supplemental Information for "The cardiac myosin binding protein-C phosphorylation state as a function of multiple protein kinase and phosphatase activities"

#### Supplemental Information Text

We examined the behaviour of the model during dynamic changes in PKA activity. Following the onset of a constant 1-hour pulse of PKA activity in the presence of constant PP1 activity, cMyBP-C phosphorylation approaches a steady-state whose phosphorylation-site distribution depends on the respective PKA and PP1 concentrations present as do the durations of the transient phase prior to reaching steady state (**Figure S21A**). Interestingly, dephosphorylation time-courses following kinase removal converged on a slowly decreasing 1P trajectory that only depends on the total PP1 concentration and results from the slow α- cMyBPC dephosphorylation kinetics. Analogous simulations with PP2A instead of PP1 show qualitatively very similar dynamics (**Figure S21B**).

The model predictions were validated by pre-incubating C1mC2 with both PKA and PP1. PKA activity was initiated by adding ATP and stopped by adding excess PKA inhibitor H-89 at defined time points. The phosphorylation time-course was resolved by Phostag™-SDS-PAGE (**Figure S21C**). Similar to the steady-state experiments (Figure 3), the experimental time-courses of the individual C1mC2 phospho-species are in good agreement with the model prediction, suggesting that the model accurately describes the phosphorylation dynamics of C1mC2 in the presence of both kinases and phosphatases.

#### Supplemental Information Figures and Figure Legends

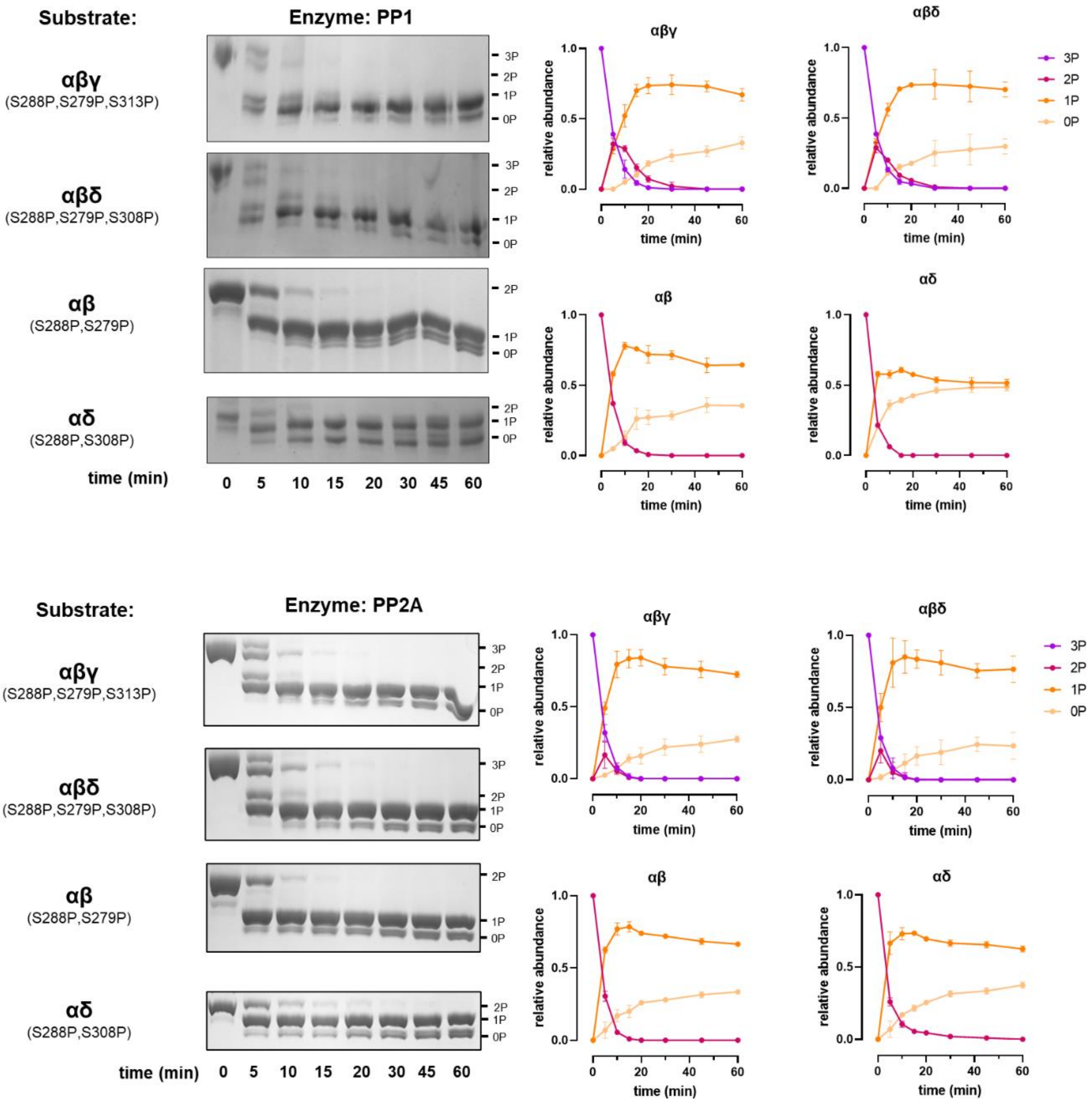


**Figure S1**: Left: additional dephosphorylation time course experiments of various C1mC2 phosphoforms by phosphatases PP1 and PP2A visualized by PhosTag$^{TM}$ SDS-PAGE gels. Right: Quantification of the data. Each datapoint represents the mean ± SD of n = 2 experiments.

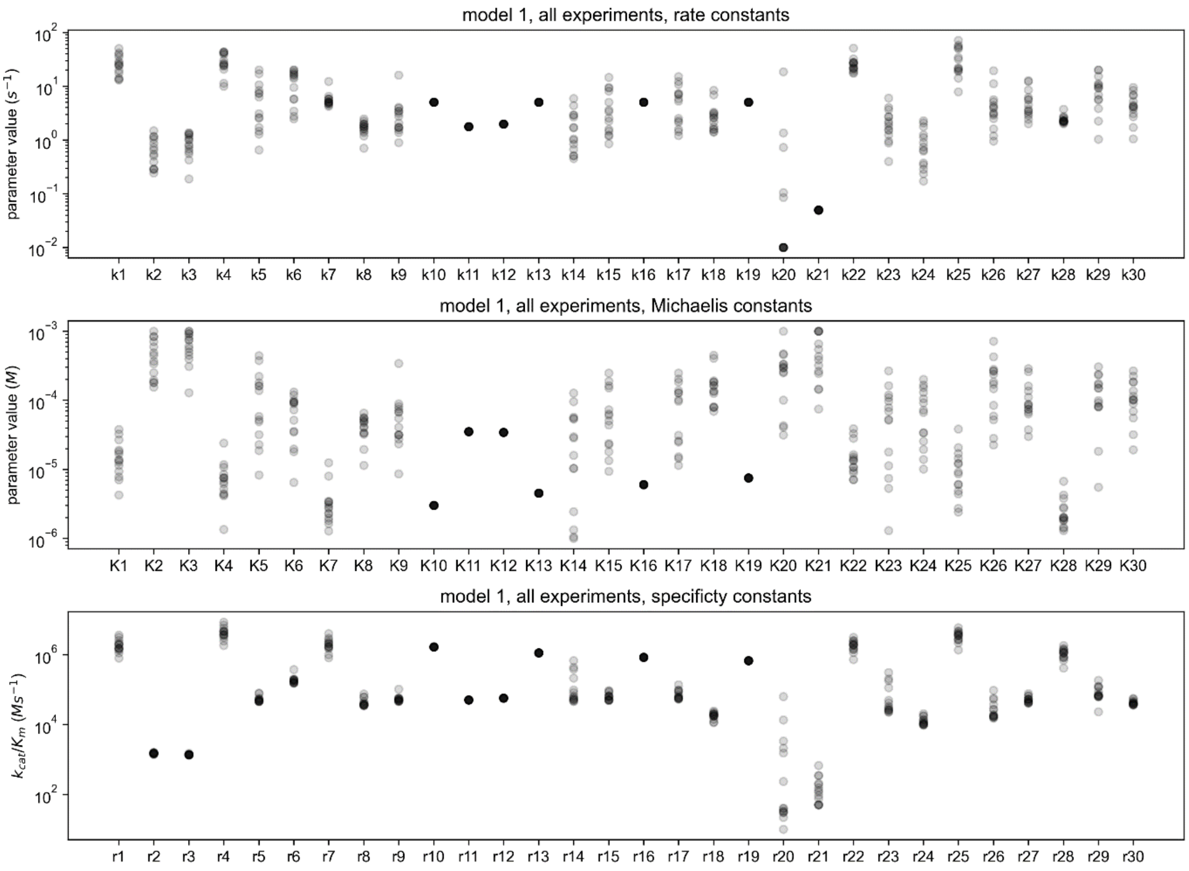


**Figure S2**: Resulting parameters for model 1 after fitting the model to all data and filtering out poorly performing parameter sets.

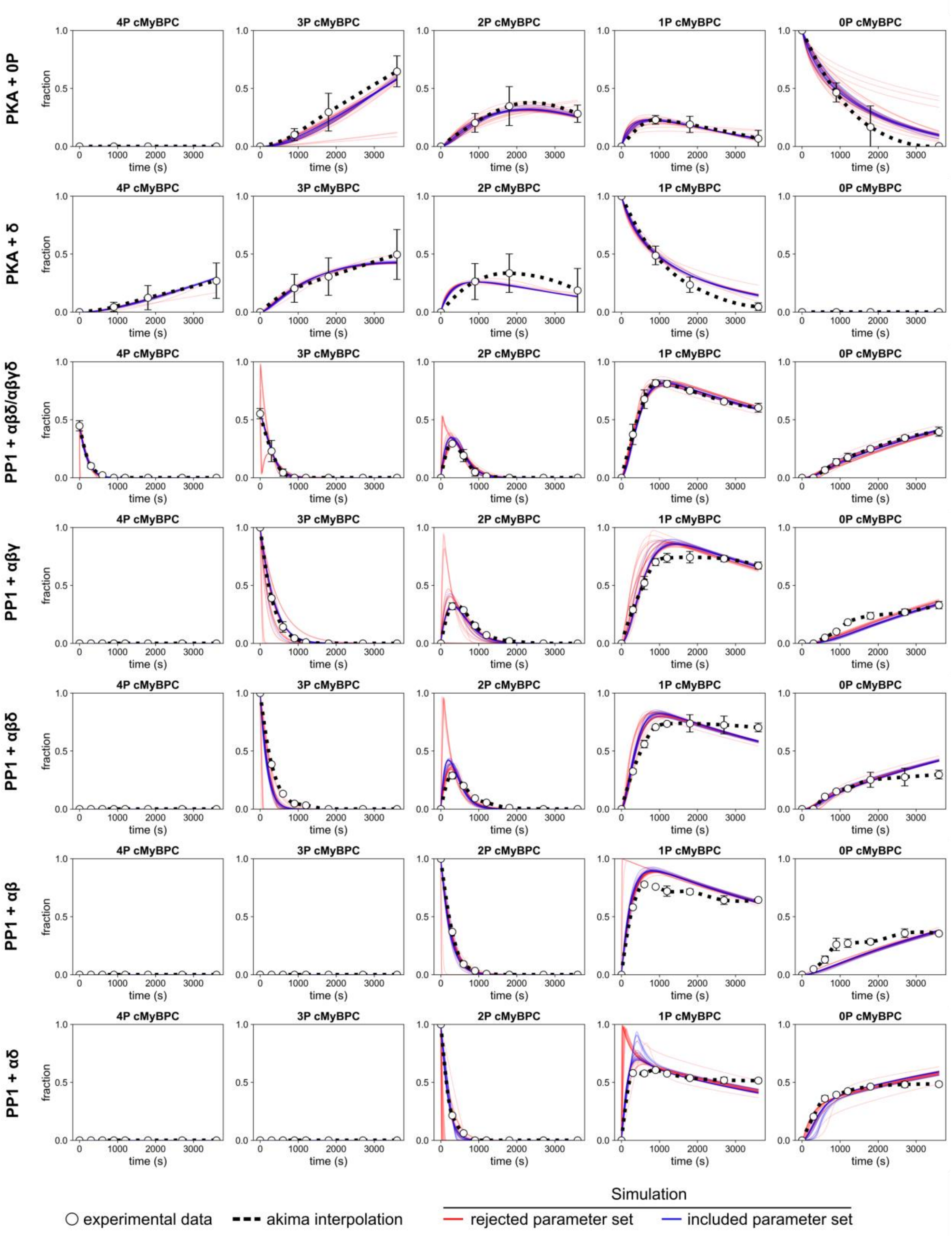


**Figure S3**: Fit of model 1 to all data, results shown for PKA and PP1 time course data. Each experimental data point represents the mean ± SD of n = 2-6 experiments.

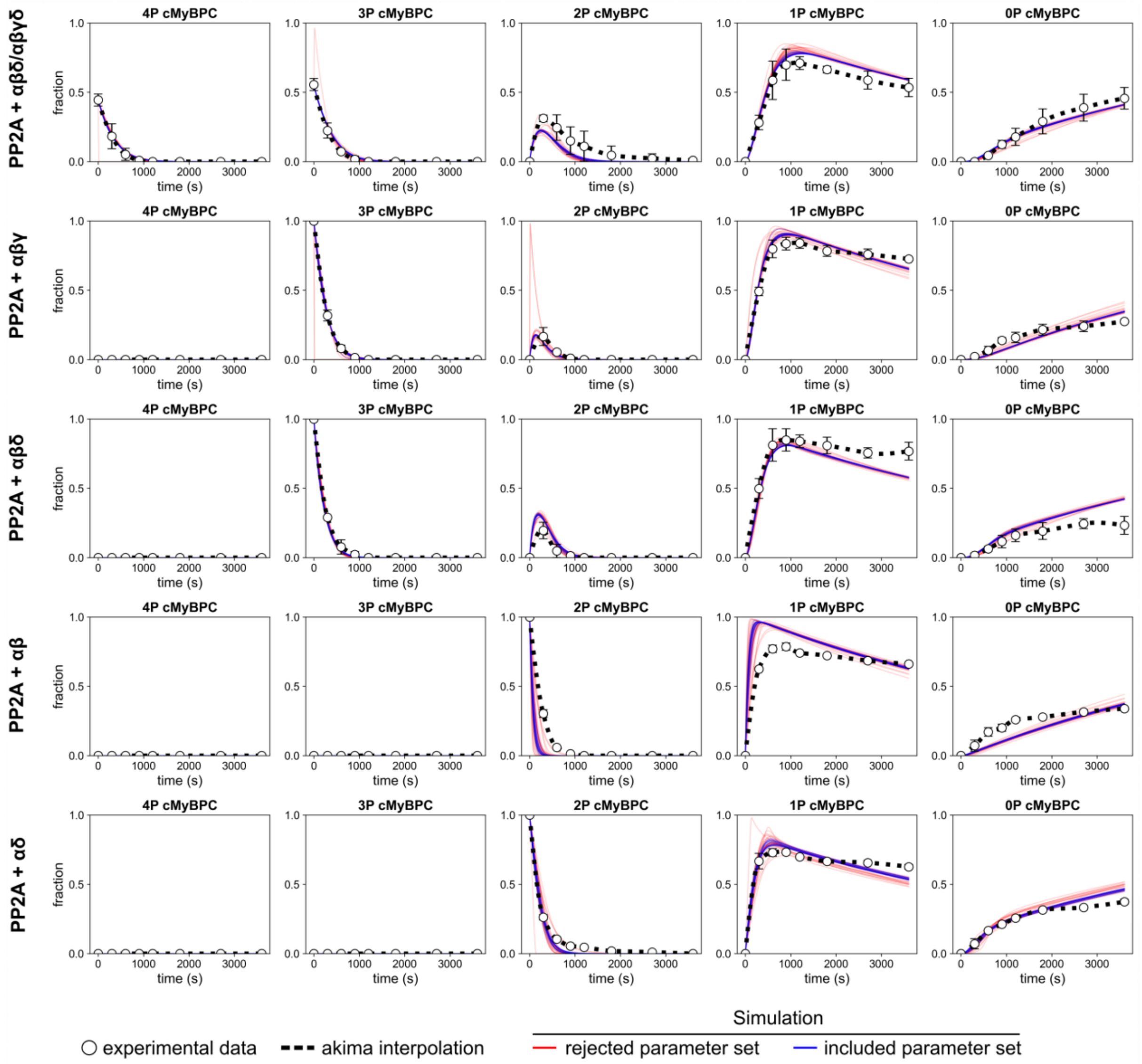


**Figure S4**: Fit of model 1 to all data, results shown for PP2A time course data. Each experimental data point represents the mean ± SD of n = 2-3 experiments.

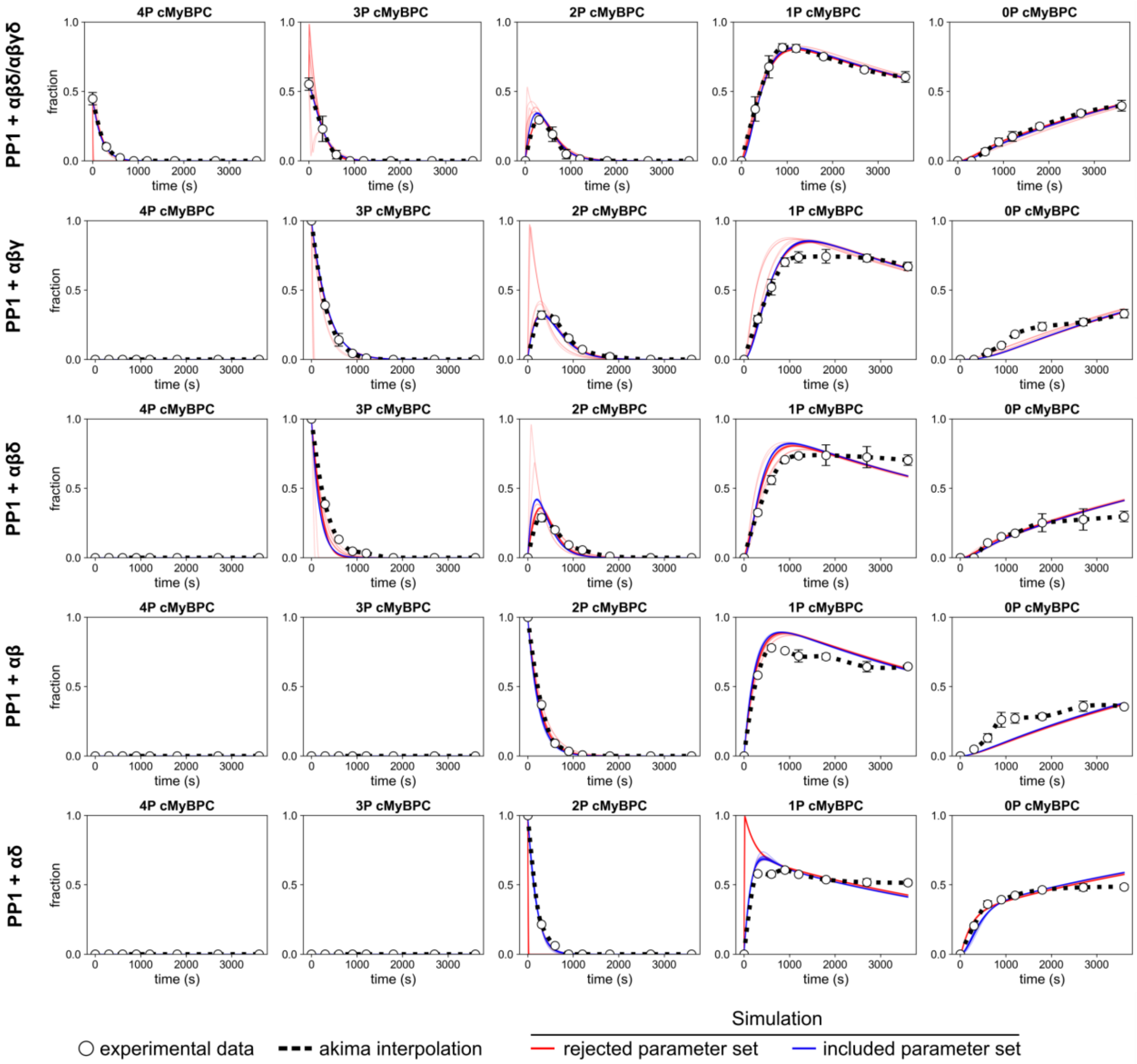


**Figure S5**: Fit of model 1 to PP1 time course data only. Each experimental data point represents the mean ± SD of n = 2-3 experiments.

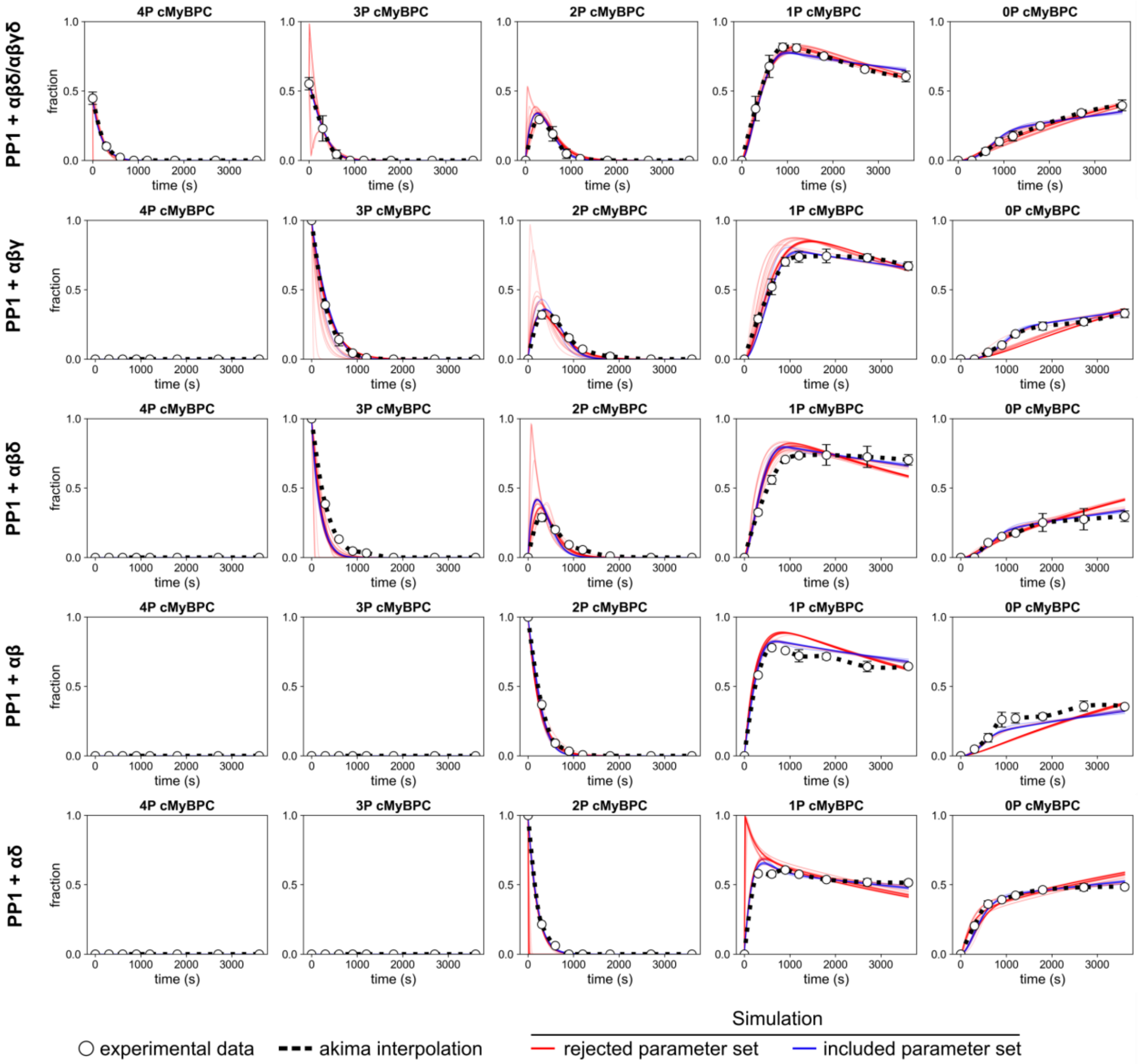


**Figure S6**: Fit of model 2 (phenomenological model of direct activation of α dephosphorylation by 2P-C1mC2) to PP1 time course data only. Each experimental data point represents the mean ± SD of n = 2-3 experiments.

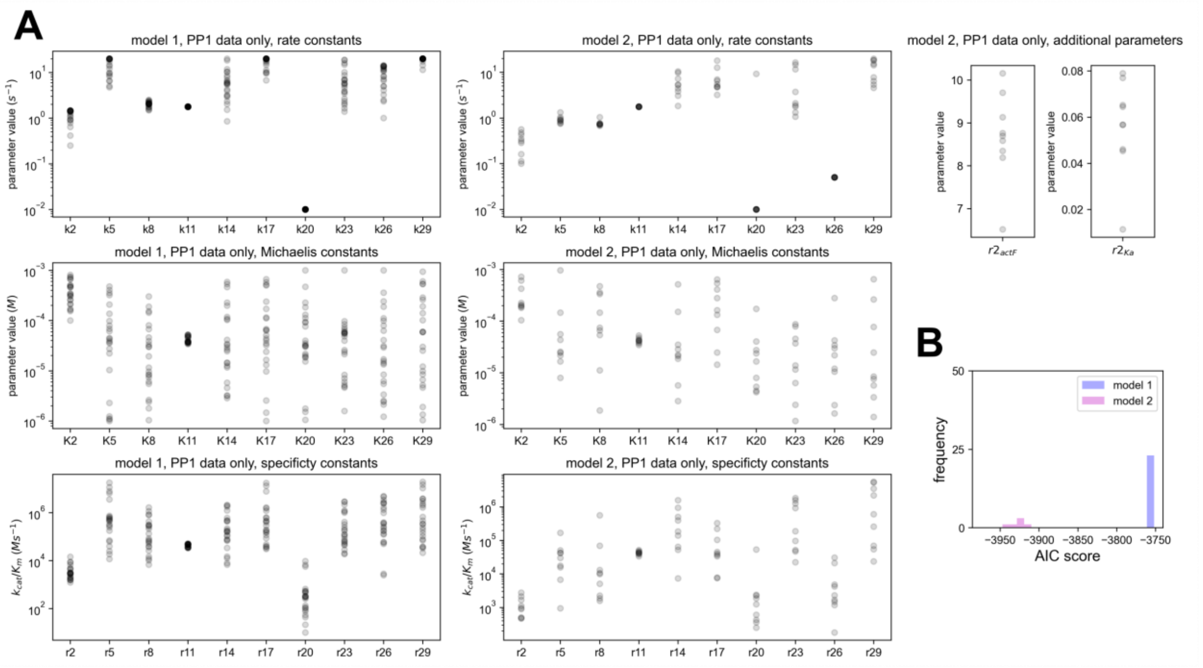


**Figure S7**: **A**, Resulting parameters for model 1 and model 2 after fitting the models to PP1 data only and filtering out poorly performing parameter sets. **B**, model comparison using the Akaike information criterion based on included parametersets after filtering (23 parameter sets included for model 1, 9 parameter sets included for model 2). Model 2 has a significantly lower AIC score than model 1 ($p = 1.6 \times 10^{-5}$; Mann-Whitney test) and thus is to be preferred over model 1.

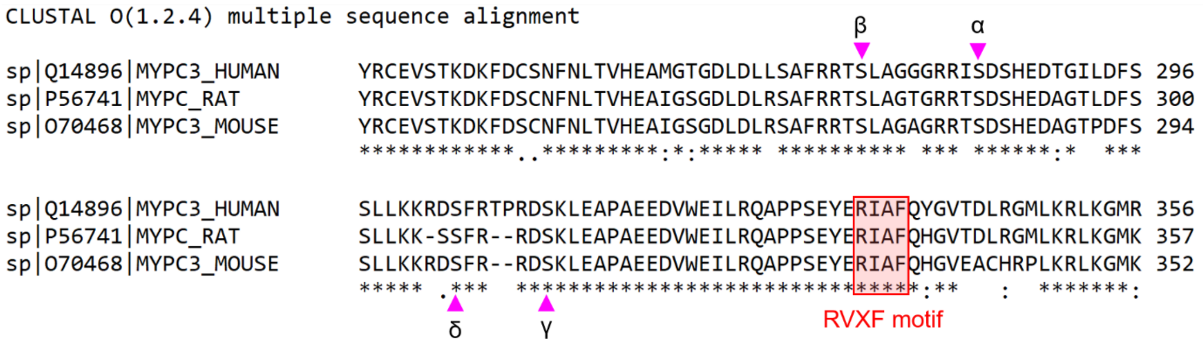


**Figure S8**: Aligned cMyBP-C sequences from human, rat and mouse. Highlighted in red is the RVXF-motif (consensus sequence: [K/R][I/V][x][F/W]) which might enable binding between cMyBP-C and PP1. Pink arrowheads indicate phosphorylation sites.

`

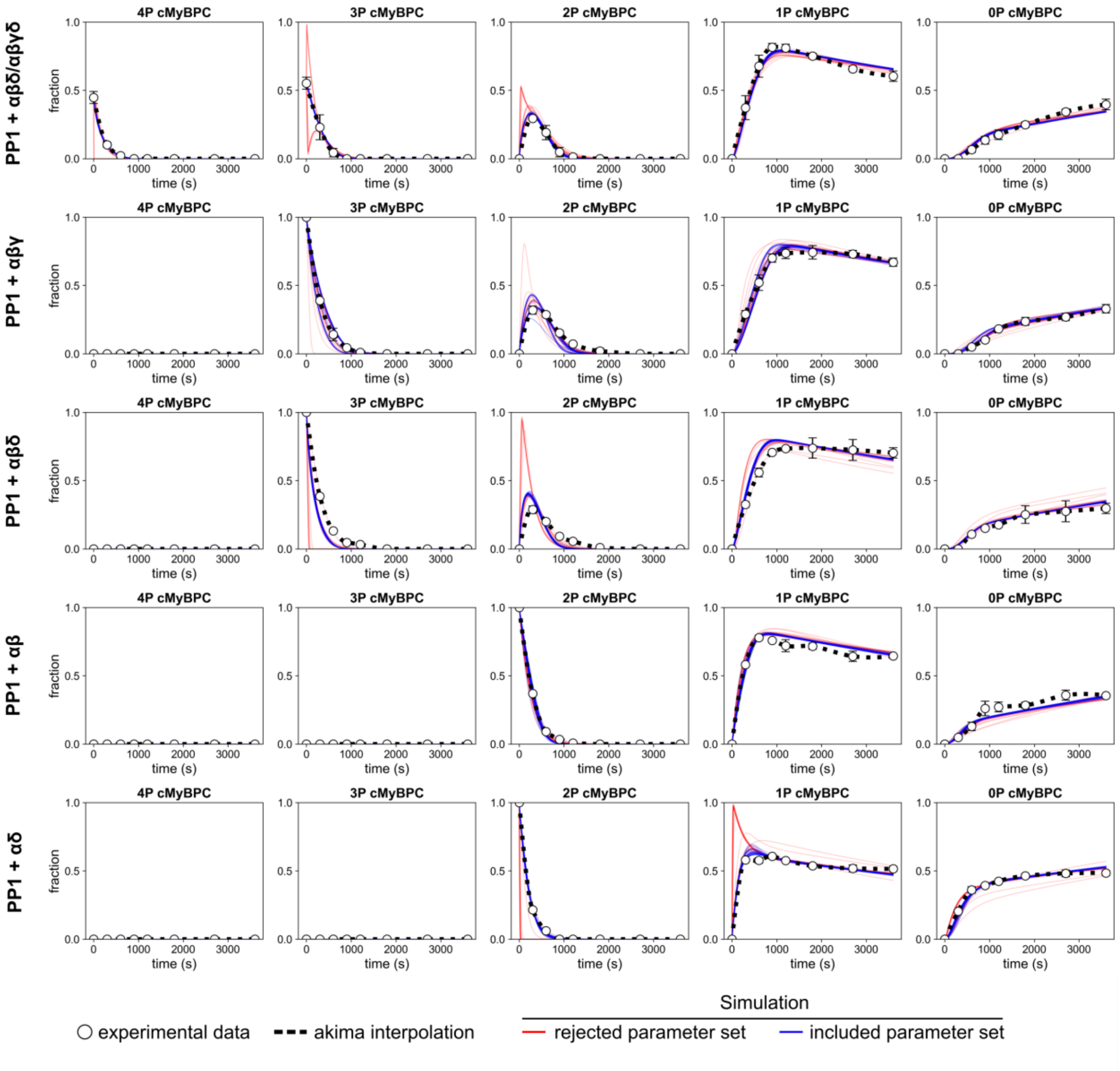


**Figure S9**: Fit of model 3 (allosteric activation of PP1) to PP1 time course data only. Each experimental data point represents the mean ± SD of n = 2-3 experiments.

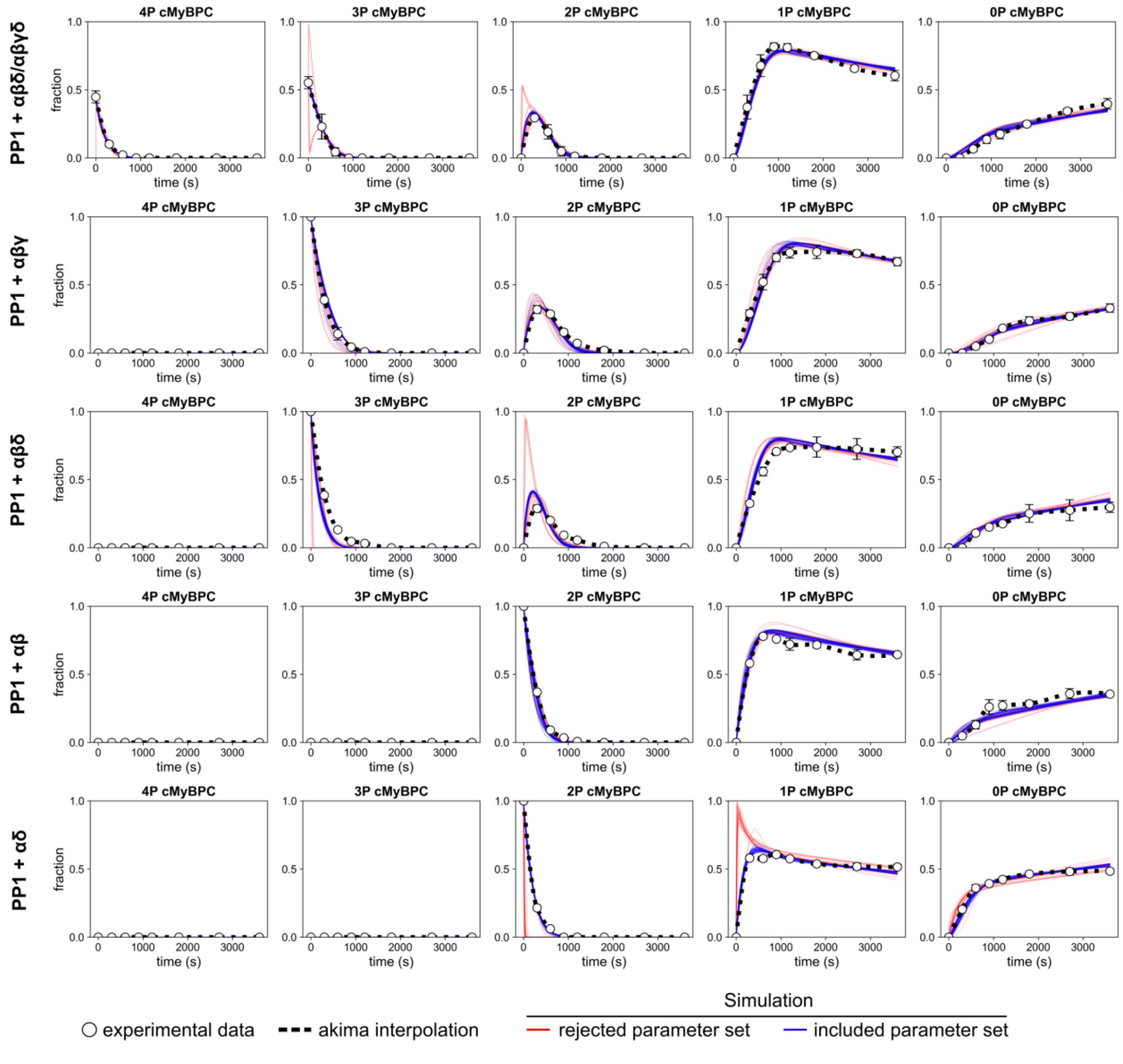


**Figure S10**: Fit of model 4 (structural transition model) to PP1 time course data only. Each experimental data point represents the mean ± SD of n = 2-3 experiments.

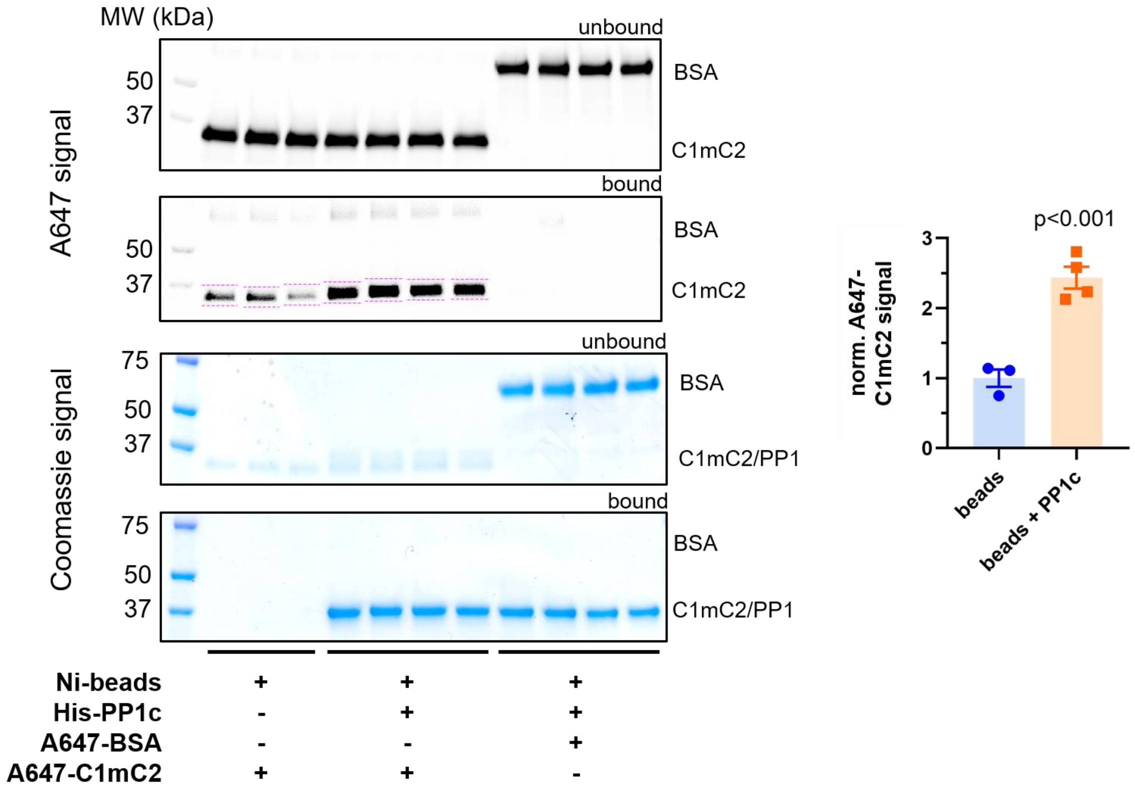


**Figure S11**: Pull-down of recombinant, Alexa-647 labelled C1mC2 domains by the recombinant, $His_6$-tagged catalytic domain of PP1. Ni-NTA beads without bait or with His-PP1c as bait were incubated with A647-C1mC2 or A647-BSA as a negative control. While some unspecific background binding between C1mC2 and beads was observed, PP1c-covered beads resulted in significantly higher C1mC2 retention (cf. quantification on the right). In contrast, BSA showed no binding to PP1.

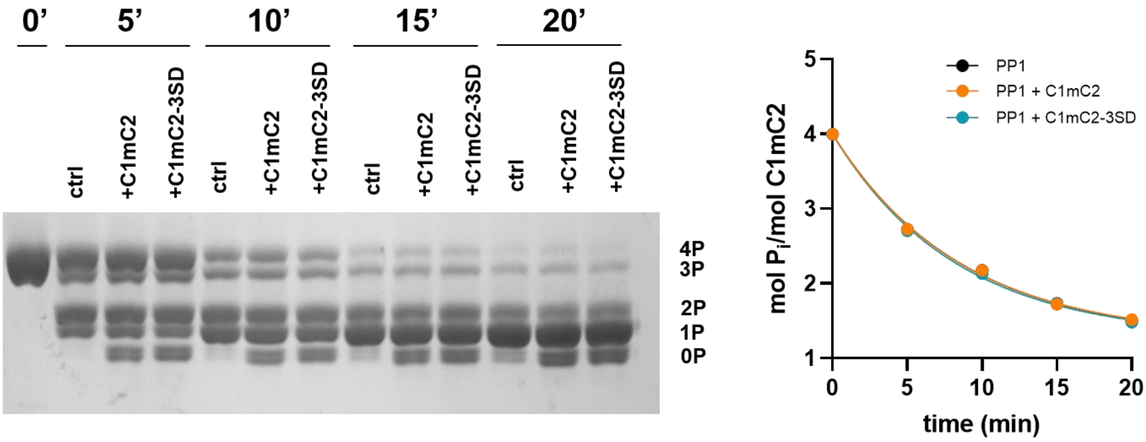


**Figure S12**: Dephosphorylation time courses for fully phosphorylated C1mC2 domain by PP1 alone, PP1 preincubated with unphosphorylated C1mC2 or PP1 preincubated with phosphomimetic Ser(279/288/313)→Asp C1mC2. No differences in the dephosphorylation rate between groups were observed.

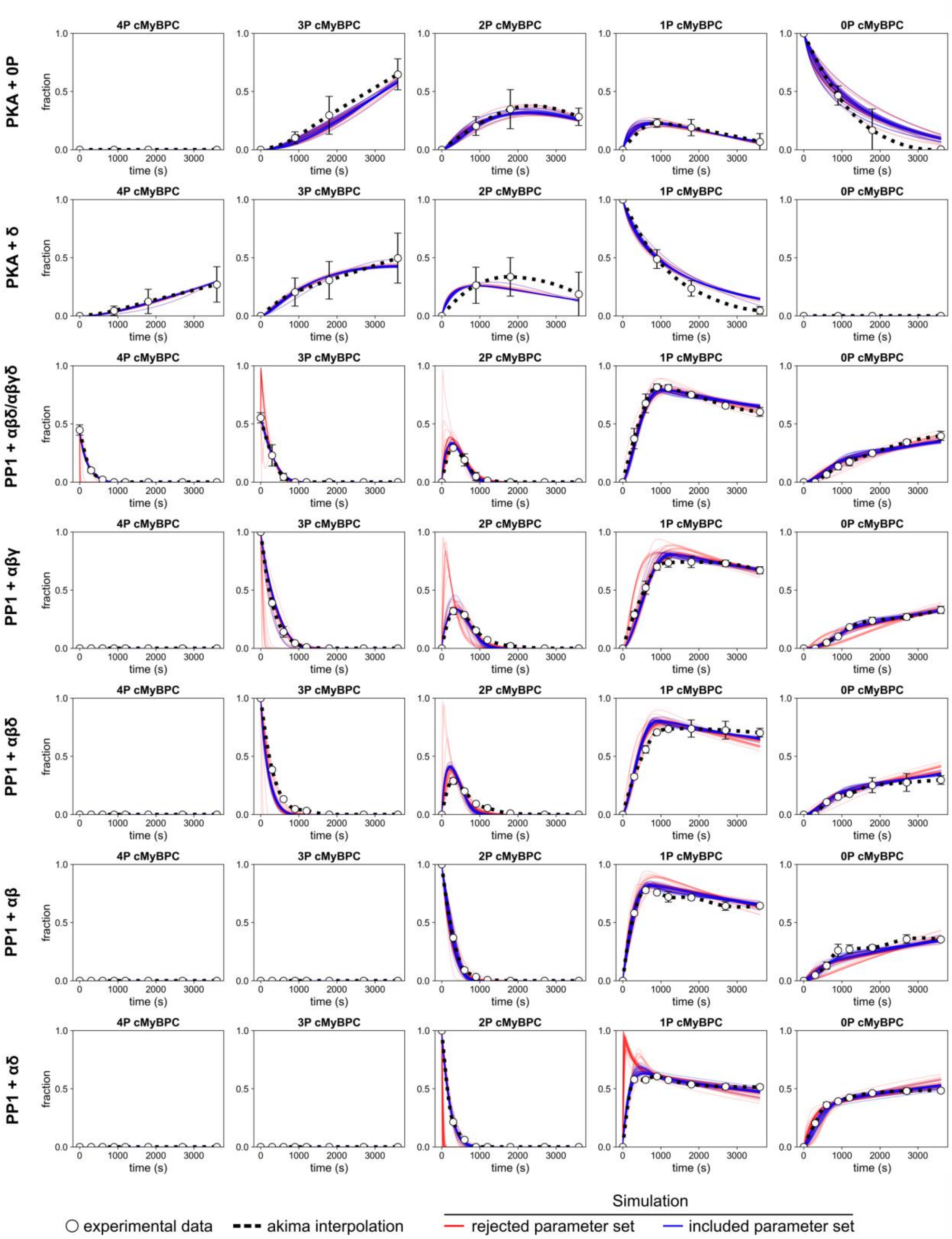


**Figure S13**: Fit of model 4 (structural transition model) to all data, results shown for PKA and PP1 time course data. Each experimental data point represents the mean ± SD of n = 2-6 experiments.

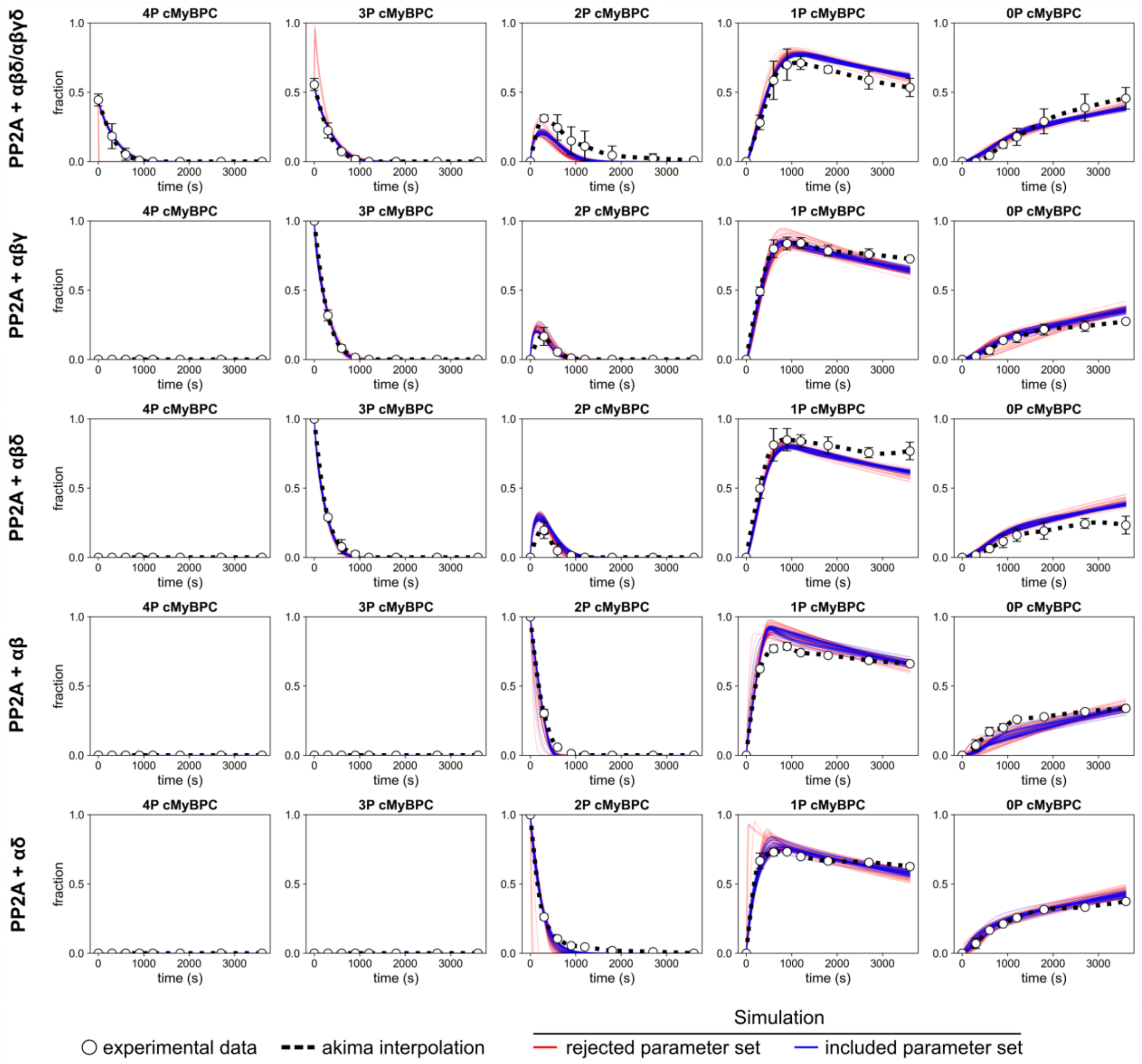


**Figure S14**: Fit of model 4 (structural transition model) to all data, results shown for PP2A time course data. Each experimental data point represents the mean ± SD of n = 2-3 experiments.

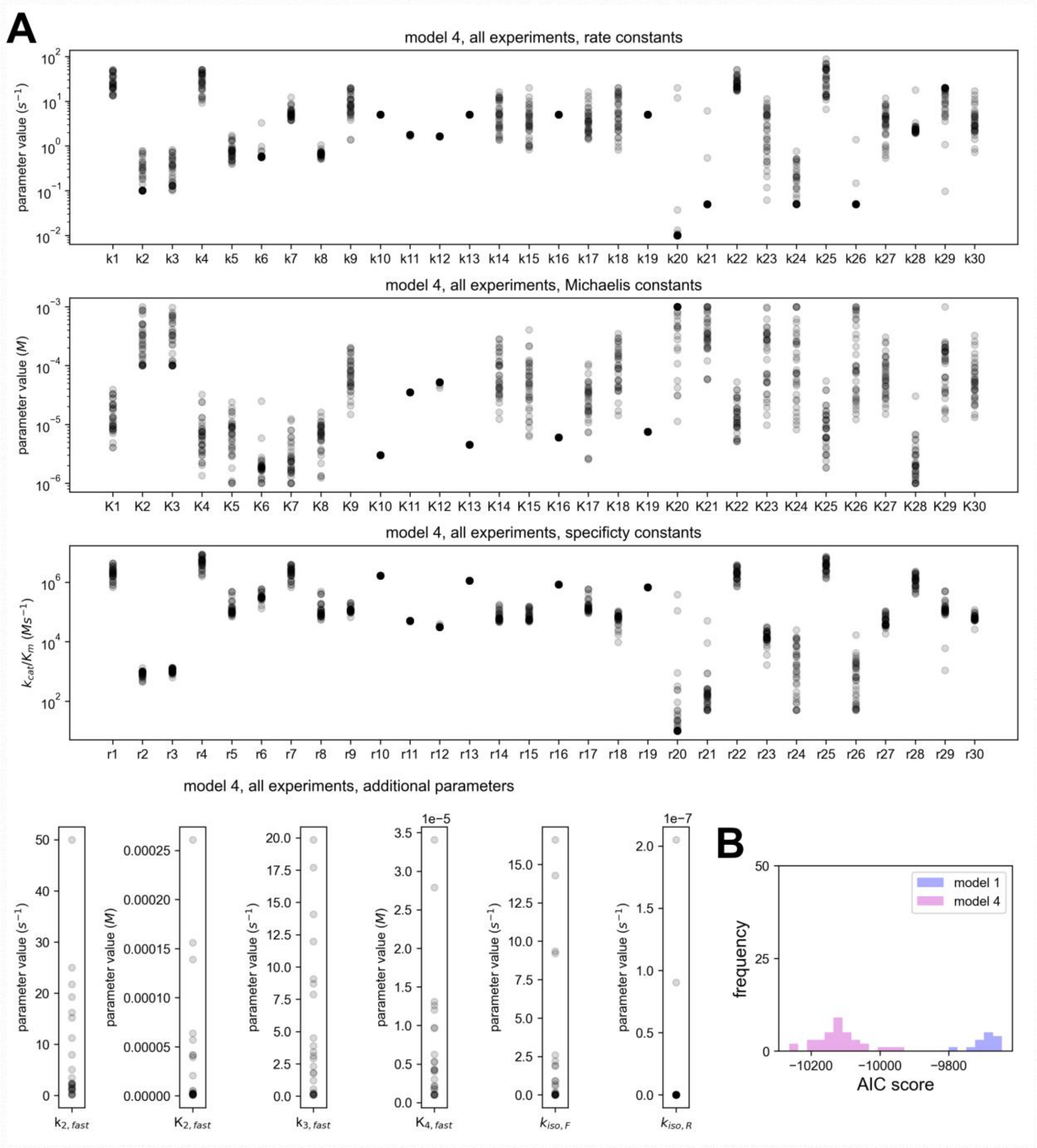


**Figure S15**: **A**, Resulting parameters for model 4 after fitting the model to all datasets and filtering out poorly performing parameter sets. **B**, Comparison of model 1 and model 4 (both fitted to all datasets) using the Akaike information criterion based on included parametersets after filtering (14 parameter sets included for model 1, 35 parameter sets included for model 4). Model 4 has a significantly lower AIC score than model 1 ($p = 7.3x10^{-30}$; Welch's t-test) and thus is to be preferred over model 1.

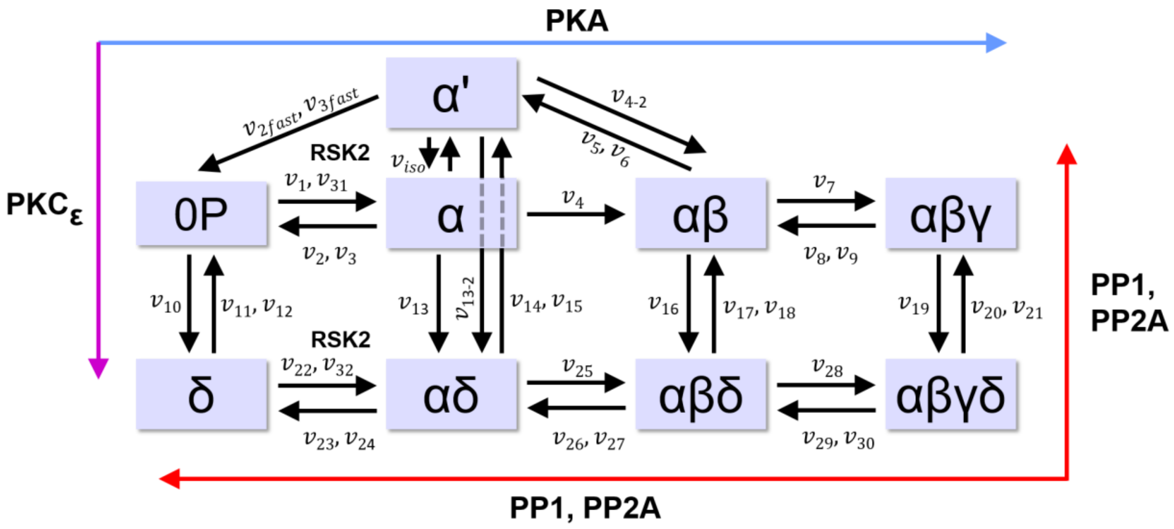


**Figure S16**: Scheme of final model (model 4 + RSK2 reactions $v_{31}$ and $v_{32}$).

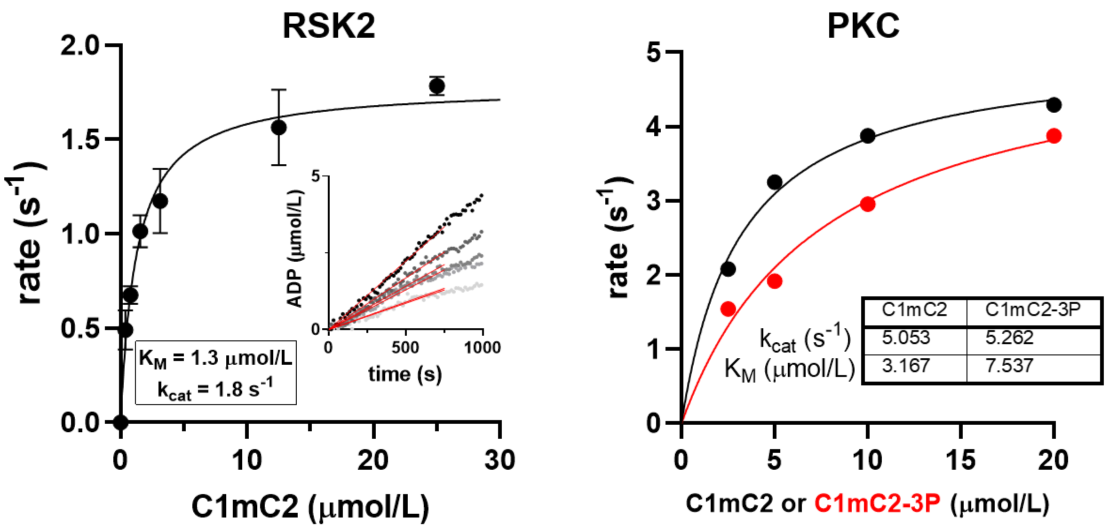


**Figure S17**: Michaelis-Menten kinetics analysis for C1mC2 phosphorylation by RSK2 (left) and PKCε (right).

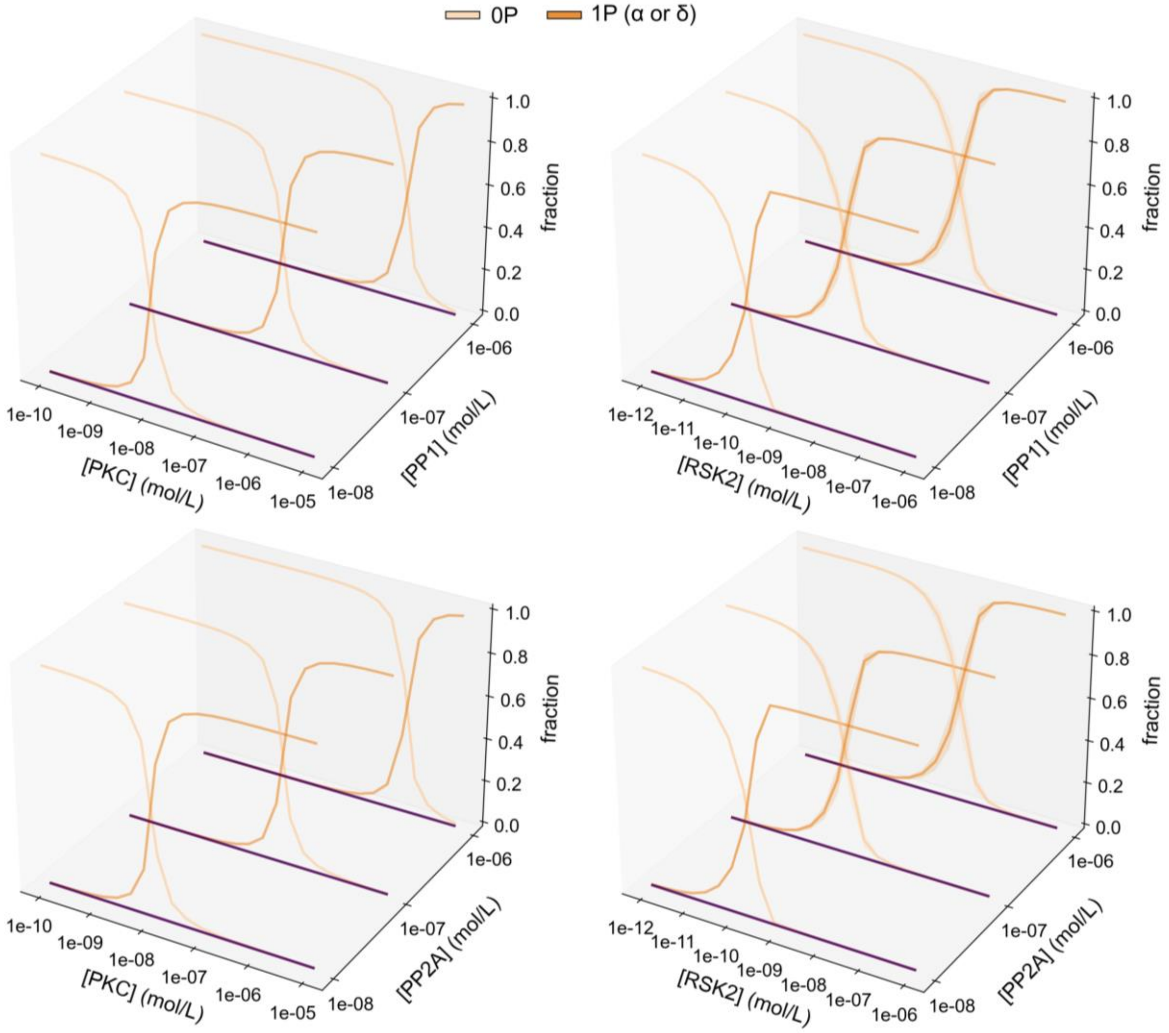


**Figure S18:** Simulated steady-state phosphorylation of cMyBP-C in the presence of PP1 or PP2A and increasing concentrations of PKC (left) or RSK2 (right).

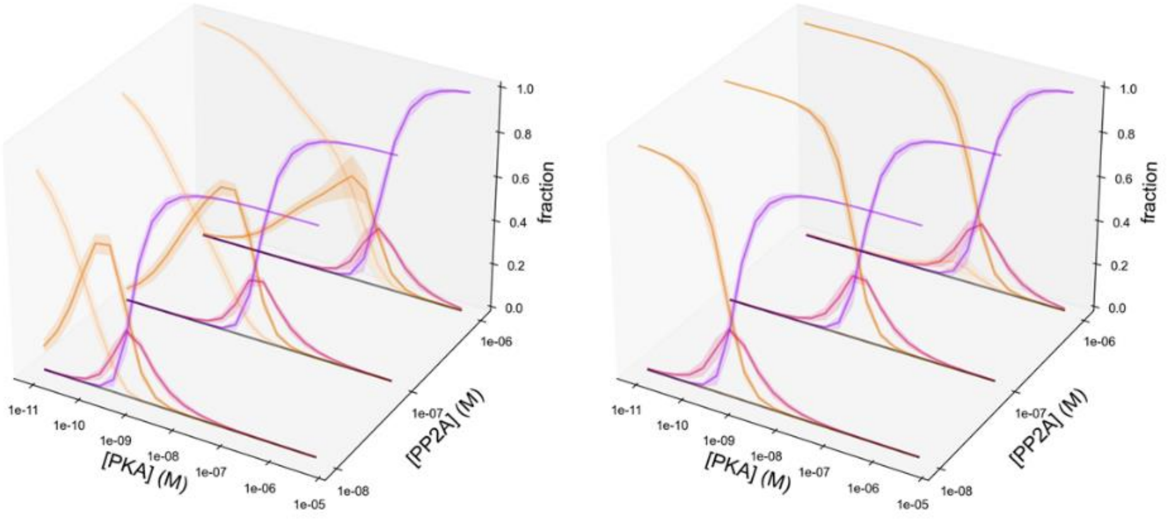


**Figure S19.** Steady-state phosphorylation simulations of cMyBP-C in the presence of PP2A and increasing concentrations of PKA (left). The effect of additional RSK2 is shown in the right panel (0P, yellow; 1P, orange; 2P, pink; 3P, purple; 4P, black).

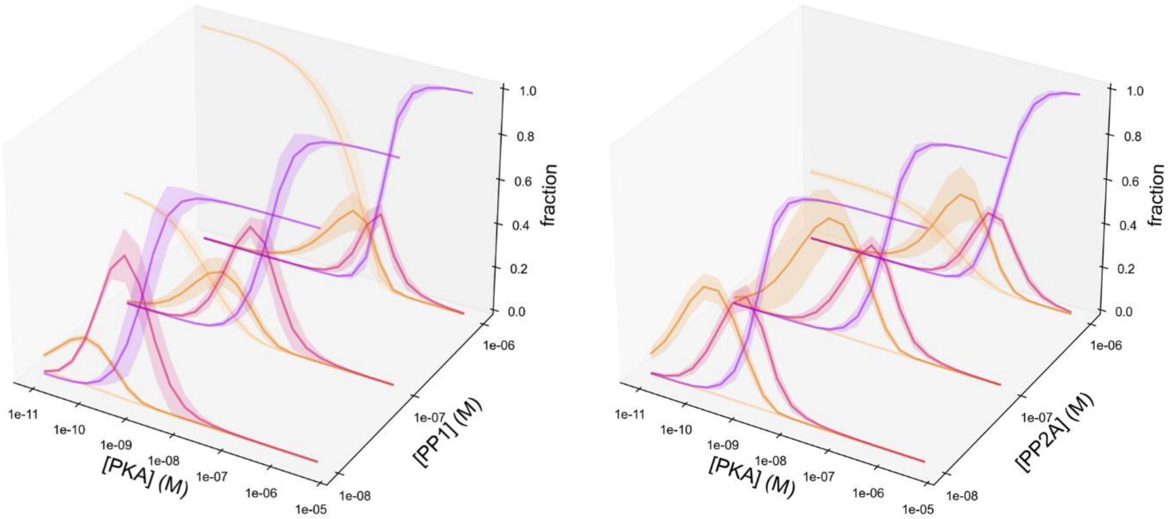


**Figure S20.** Steady-state phosphorylation simulations of cMyBP-C in the presence of PKCε and PP1 (left) or PP2A (right), and increasing concentrations of PKA with the 3P and 4P species lumped together (0P, yellow; 1P, orange; 2P, pink; 3P+4P, purple).

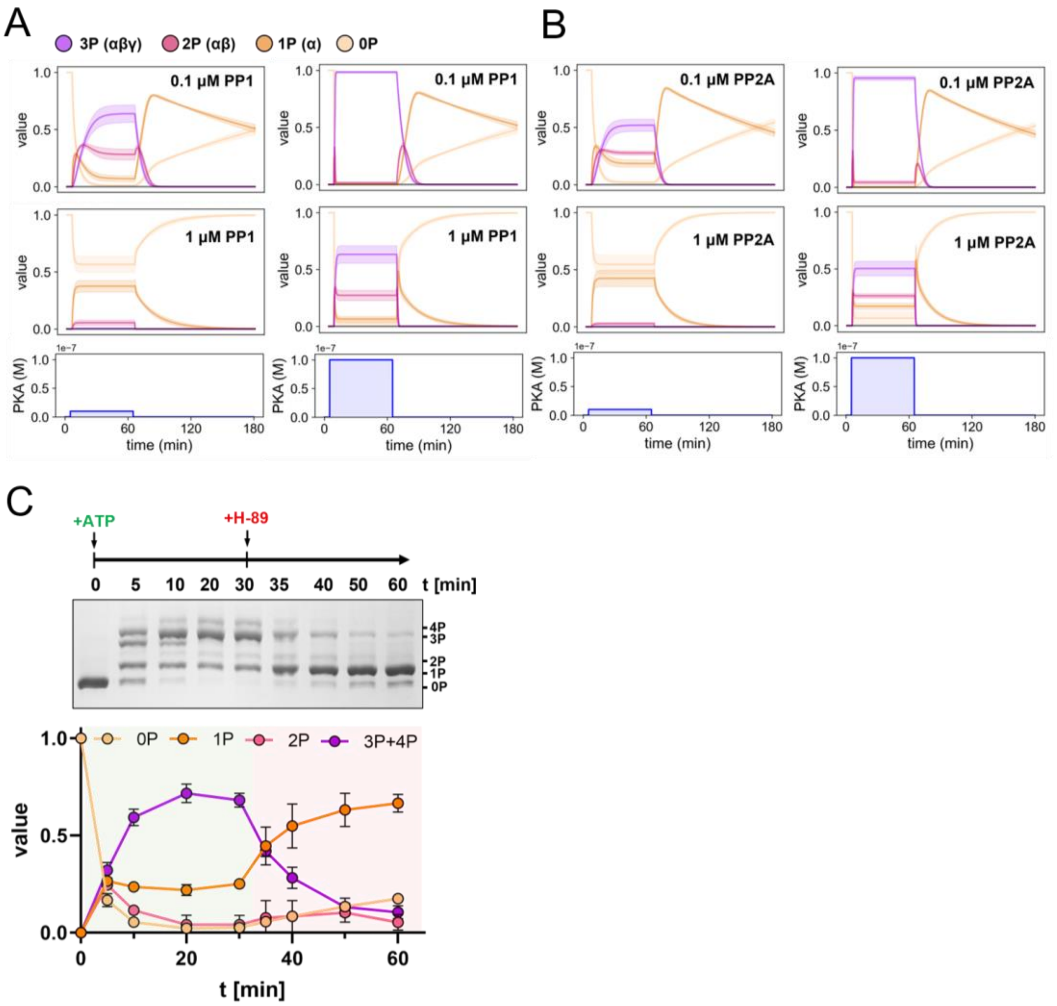


**Figure S21:** Model prediction for the time-dependent phosphorylation of cMyBP-C by PKA in the presence of (A) low and high PP1 concentration, and (B) low and high PP2A concentrations. (C) Experimental validation of the predicted time-courses using ATP and H-89 to start (t = 0 min) and stop (t = 30 min) PKA activity at defined time points, respectively.

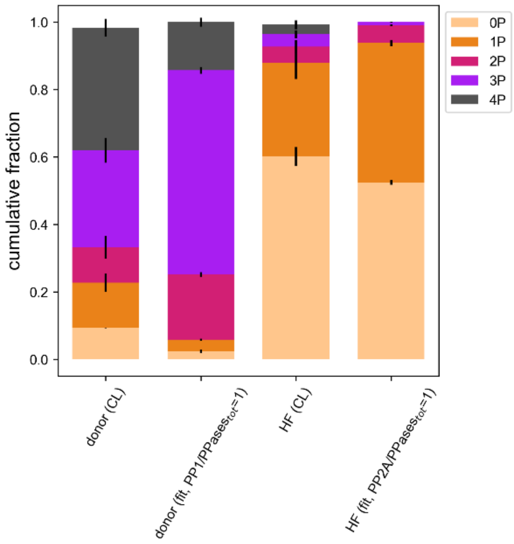


**Figure S22:** Control experiment with clamped phosphatase ratios. Like in Figure 4A (main text), the model was tested for consistency with experimental data on cMyBP-C basal phosphorylation states in hearts from healthy donors or HF patients reported in Copeland *et al.* 2010 (CL) by fitting the model using only the enzyme concentrations as free parameters (fit). However, for donor heart data, the PP1/PPases$_{tot}$ was set to 1 (i.e. no PP2A was present), whereas for HF data, the PP2A/PPases$_{tot}$ was fixed to 1 (i.e. no PP1 was present).

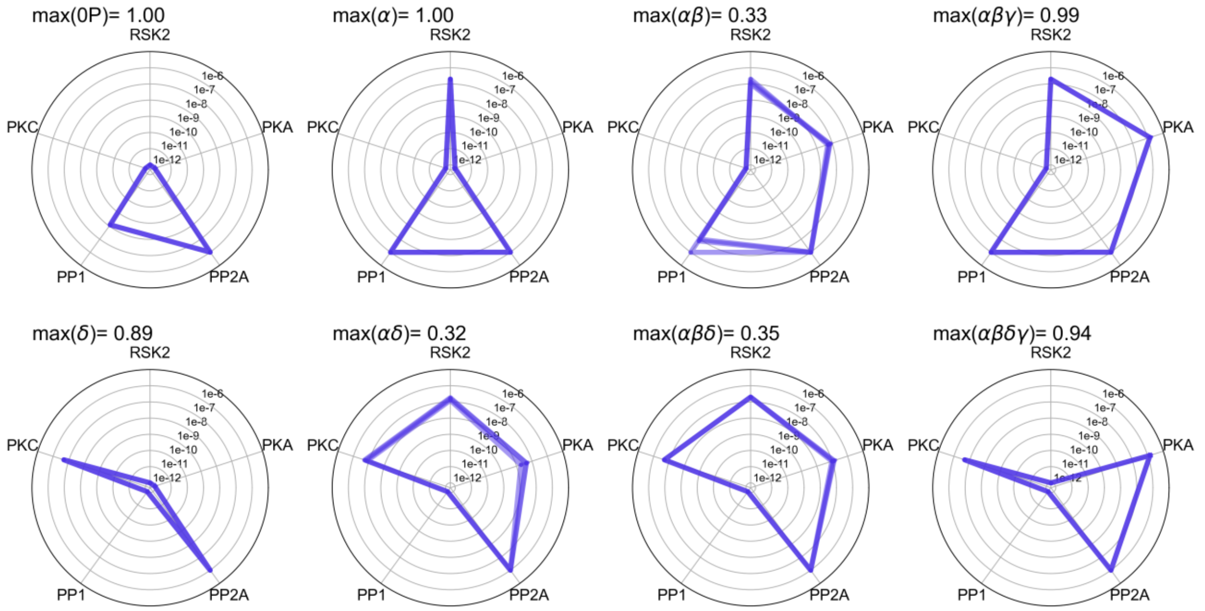


**Figure S23**: Optimization of cMyBP-C states under physiological conditions. Spider plots show enzyme vectors at which the respective cMyBP-C phosphorylation is at its maximally possible fraction.

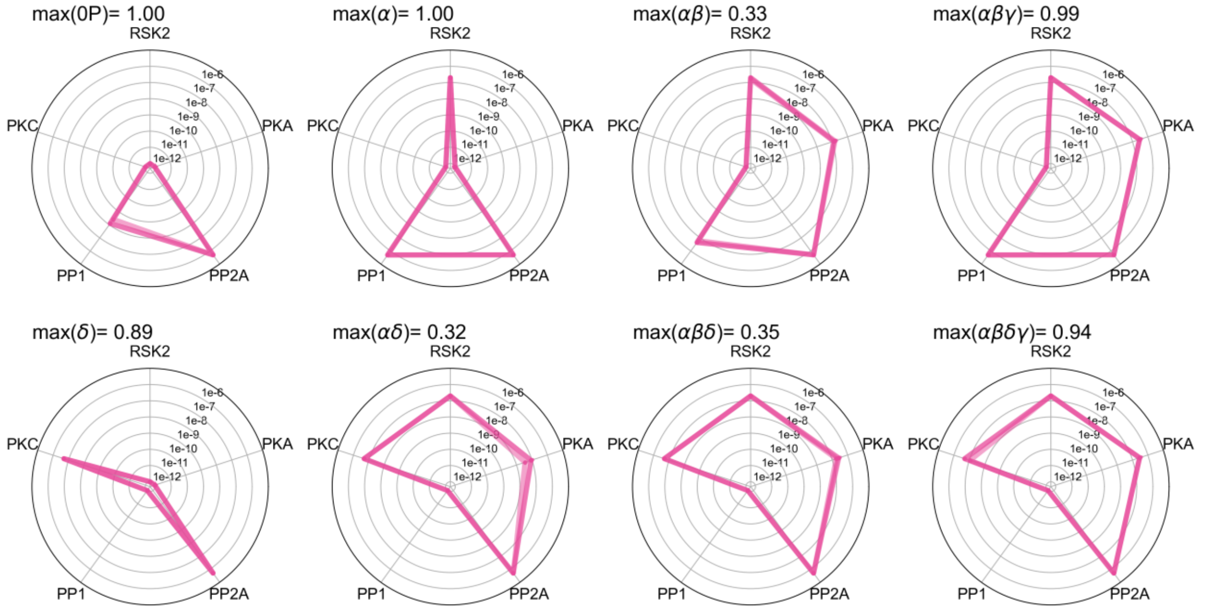


**Figure S24**: Optimization of cMyBP-C states under HF conditions. Spider plots show enzyme vectors at which the respective cMyBP-C phosphorylation is at its maximally possible fraction.

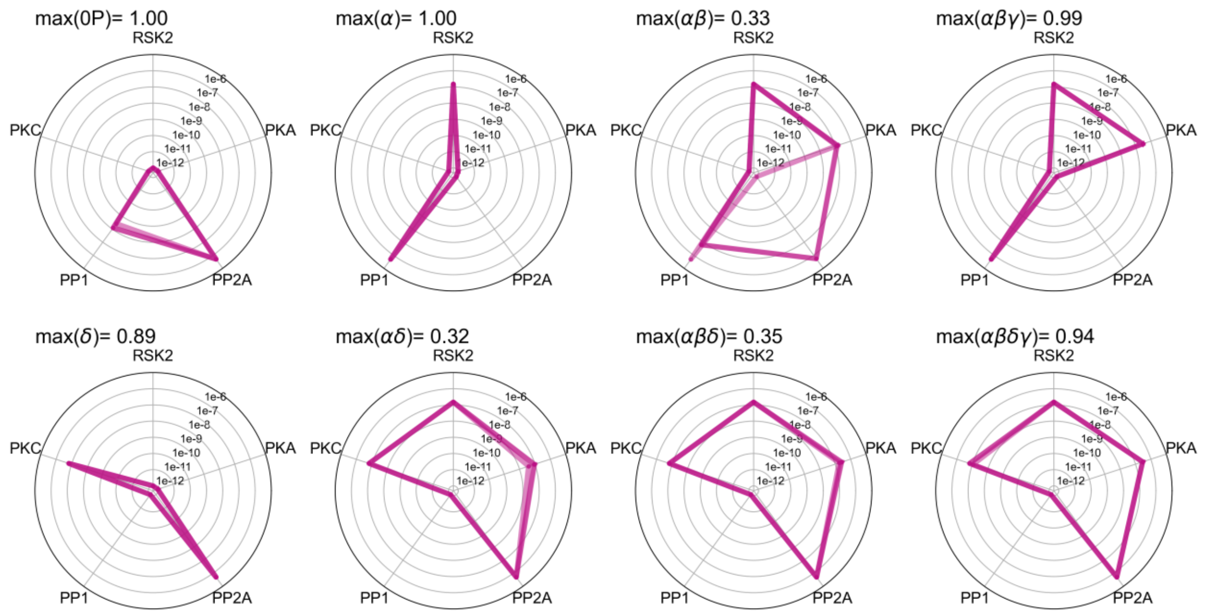


**Figure S25**: Optimization of cMyBP-C states under HF conditions with additional restriction of PKC and RSK2 concentrations. Spider plots show enzyme vectors at which the respective cMyBP-C phosphorylation is at its maximally possible fraction.

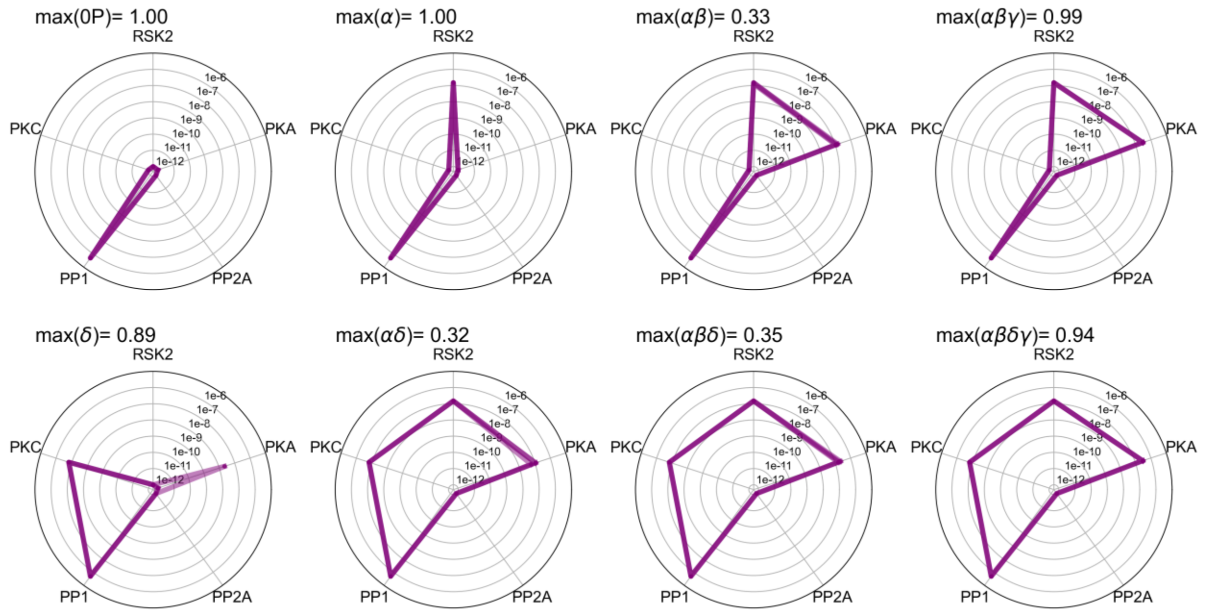


**Figure S26**: Optimization of cMyBP-C states under HF conditions with additional restriction of PKC and RSK2 concentrations and clamped PP1. Spider plots show enzyme vectors at which the respective cMyBP-C phosphorylation is at its maximally possible fraction.

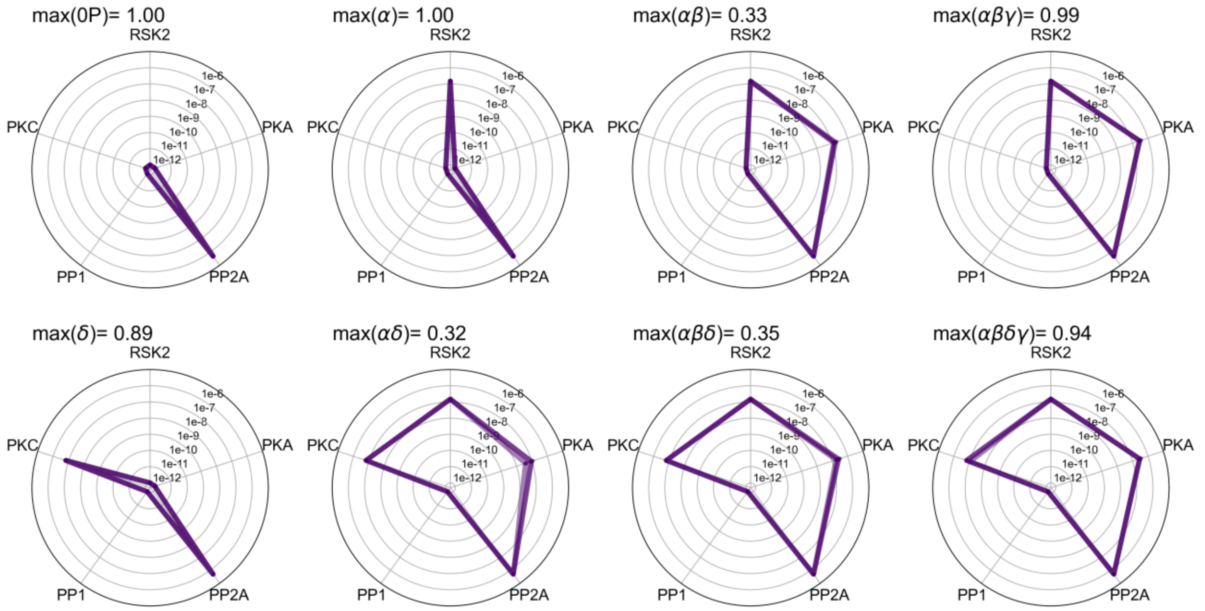


**Figure S27**: Optimization of cMyBP-C states under HF conditions with additional restriction of PKC and RSK2 concentrations and clamped PP2A. Spider plots show enzyme vectors at which the respective cMyBP-C phosphorylation is at its maximally possible fraction.

Table S1: Statistical comparisons of Hill-coefficients $n_H$ and EC50 values

| **Dose response simulations** | **Comparisons** | **Test** | **p-value** | **Significant?** |
|---|---|---|---|---|
| PKA vs PP1 | $n_H$ at different [Phosphatase] ($10^{-8}$ vs $10^{-7}$ vs $10^{-6}$ mol/L) | Kruskal-Wallis | $2.08 \times 10^{-7}$ | Yes |
| PKA + 100 nmol/L RSK2 vs PP1 | $n_H$ at different [Phosphatase] ($10^{-8}$ vs $10^{-7}$ vs $10^{-6}$ mol/L) | Kruskal-Wallis | $2.20 \times 10^{-9}$ | Yes |
| PKA + 100 nmol/L PKC vs PP1 | $n_H$ at different [Phosphatase] ($10^{-8}$ vs $10^{-7}$ vs $10^{-6}$ mol/L) | Kruskal-Wallis | $1.27 \times 10^{-18}$ | Yes |
| PKA vs PP2A | $n_H$ at different [Phosphatase] ($10^{-8}$ vs $10^{-7}$ vs $10^{-6}$ mol/L) | Kruskal-Wallis | $2.31 \times 10^{-7}$ | Yes |
| PKA + 100 nmol/L RSK2 vs PP2A | $n_H$ at different [Phosphatase] ($10^{-8}$ vs $10^{-7}$ vs $10^{-6}$ mol/L) | Kruskal-Wallis | $2.28 \times 10^{-5}$ | Yes |
| PKA + 100 nmol/L PKC vs PP2A | $n_H$ at different [Phosphatase] ($10^{-8}$ vs $10^{-7}$ vs $10^{-6}$ mol/L) | Kruskal-Wallis | $1.82 \times 10^{-13}$ | Yes |
| PKA vs PP1, PKA + 100 nmol/L RSK2 vs PP1, PKA + 100 nmol/L PKC vs PP1 | $n_H$ across dose responses at [Phosphatase] = $10^{-8}$ mol/L | Kruskal-Wallis | $3.23 \times 10^{-15}$ | Yes |
| PKA vs PP1, PKA + 100 nmol/L RSK2 vs PP1, PKA + 100 nmol/L PKC vs PP1 | $n_H$ across dose responses at [Phosphatase] = $10^{-7}$ mol/L | Kruskal-Wallis | $3.23 \times 10^{-15}$ | Yes |
| PKA vs PP1, PKA + 100 nmol/L RSK2 vs PP1, PKA + 100 nmol/L PKC vs PP1 | $n_H$ across dose responses at [Phosphatase] = $10^{-6}$ mol/L | Kruskal-Wallis | 0.07 | No |
| PKA vs PP2A, PKA + 100 nmol/L RSK2 vs PP2A, PKA + 100 nmol/L PKC vs PP2A | $n_H$ across dose responses at [Phosphatase] = $10^{-8}$ mol/L | Kruskal-Wallis | $1.04 \times 10^{-6}$ | Yes |
| PKA vs PP2A, PKA + 100 nmol/L RSK2 vs PP2A, PKA + 100 nmol/L PKC vs PP2A | $n_H$ across dose responses at [Phosphatase] = $10^{-7}$ mol/L | Kruskal-Wallis | 0.065 | No |
| PKA vs PP2A, PKA + 100 nmol/L RSK2 vs PP2A, PKA + 100 nmol/L PKC vs PP2A | $n_H$ across dose responses at [Phosphatase] = $10^{-6}$ mol/L | Kruskal-Wallis | 0.0016 | Yes |
| PKA vs PP1, PKA vs PP2A | $n_H$ for PP1 vs PP2A at [Phosphatase] = $10^{-8}$ mol/L | Mann-Whitney | $1.88 \times 10^{-7}$ | Yes |
| PKA vs PP1, PKA vs PP2A | $n_H$ for PP1 vs PP2A at [Phosphatase] = $10^{-7}$ mol/L | Mann-Whitney | $2.51 \times 10^{-9}$ | Yes |
| PKA vs PP1, PKA vs PP2A | $n_H$ for PP1 vs PP2A at [Phosphatase] = $10^{-6}$ mol/L | Mann-Whitney | $4.70 \times 10^{-9}$ | Yes |
| PKA + 100 nmol/L RSK2 vs PP1, PKA + 100 nmol/L RSK2 vs PP2A | $n_H$ for PP1 vs PP2A at [Phosphatase] = $10^{-8}$ mol/L | Mann-Whitney | $5.06 \times 10^{-5}$ | Yes |
| PKA + 100 nmol/L RSK2 vs PP1, PKA + 100 nmol/L RSK2 vs PP2A | $n_H$ for PP1 vs PP2A at [Phosphatase] = $10^{-7}$ mol/L | Mann-Whitney | $5.06 \times 10^{-9}$ | Yes |
| PKA + 100 nmol/L RSK2 vs PP1, PKA + 100 nmol/L RSK2 vs PP2A | $n_H$ for PP1 vs PP2A at [Phosphatase] = $10^{-6}$ mol/L | Mann-Whitney | $1.45 \times 10^{-8}$ | Yes |
| PKA + 100 nmol/L PKC vs PP1, PKA + 100 nmol/L PKC vs PP2A | $n_H$ for PP1 vs PP2A at [Phosphatase] = $10^{-8}$ mol/L | Mann-Whitney | 0.6 | No |
| PKA + 100 nmol/L PKC vs PP1, PKA + 100 nmol/L PKC vs PP2A | $n_H$ for PP1 vs PP2A at [Phosphatase] = $10^{-7}$ mol/L | Mann-Whitney | 0.5 | No |
| PKA + 100 nmol/L PKC vs PP1, PKA + 100 nmol/L PKC vs PP2A | $n_H$ for PP1 vs PP2A at [Phosphatase] = $10^{-6}$ mol/L | Mann-Whitney | $3.97 \times 10^{-10}$ | Yes |
| PKA vs PP1, PKA vs PP2A | EC50 for PP1 vs PP2A at [Phosphatase] = $10^{-8}$ mol/L | Mann-Whitney | $1.12 \times 10^{-5}$ | Yes |
| PKA vs PP1, PKA vs PP2A | EC50 for PP1 vs PP2A at [Phosphatase] = $10^{-7}$ mol/L | Mann-Whitney | $1.37 \times 10^{-8}$ | Yes |
| PKA vs PP1, PKA vs PP2A | EC50 for PP1 vs PP2A at [Phosphatase] = $10^{-6}$ mol/L | Mann-Whitney | $1.37 \times 10^{-8}$ | Yes |
| PKA + 100 nmol/L RSK2 vs PP1, PKA + 100 nmol/L RSK2 vs PP2A | EC50 for PP1 vs PP2A at [Phosphatase] = $10^{-8}$ mol/L | Mann-Whitney | $8.67 \times 10^{-7}$ | Yes |
| PKA + 100 nmol/L RSK2 vs PP1, PKA + 100 nmol/L RSK2 vs PP2A | EC50 for PP1 vs PP2A at [Phosphatase] = $10^{-7}$ mol/L | Mann-Whitney | $2.05 \times 10^{-8}$ | Yes |
| PKA + 100 nmol/L RSK2 vs PP1, PKA + 100 nmol/L RSK2 vs PP2A | EC50 for PP1 vs PP2A at [Phosphatase] = $10^{-6}$ mol/L | Mann-Whitney | $1.37 \times 10^{-8}$ | Yes |
| PKA + 100 nmol/L PKC vs PP1, PKA + 100 nmol/L PKC vs PP2A | EC50 for PP1 vs PP2A at [Phosphatase] = $10^{-8}$ mol/L | Mann-Whitney | 0.0002 | Yes |
| PKA + 100 nmol/L PKC vs PP1, PKA + 100 nmol/L PKC vs PP2A | EC50 for PP1 vs PP2A at [Phosphatase] = $10^{-7}$ mol/L | Mann-Whitney | $8.61 \times 10^{-8}$ | Yes |
| PKA + 100 nmol/L PKC vs PP1, PKA + 100 nmol/L PKC vs PP2A | EC50 for PP1 vs PP2A at [Phosphatase] = $10^{-6}$ mol/L | Mann-Whitney | $6.74 \times 10^{-11}$ | Yes |

### Appendix: model equations

**Model 1** (Michaelis-Menten-type reactions only)

*Substrate competition terms*

$$\kappa_{PKA} = \frac{0P}{K_1} + \frac{\alpha}{K_4} + \frac{\alpha\beta}{K_7} + \frac{\delta}{K_{22}} + \frac{\alpha\delta}{K_{25}} + \frac{\alpha\beta\delta}{K_{28}} \qquad \kappa_{PKC} = \frac{0P}{K_{10}} + \frac{\alpha}{K_{13}} + \frac{\alpha\beta}{K_{16}} + \frac{\alpha\beta\gamma}{K_{19}}$$

$$\kappa_{PP1} = \frac{\alpha}{K_2} + \frac{\alpha\beta}{K_5} + \frac{\alpha\beta\gamma}{K_8} + \frac{\delta}{K_{11}} + \frac{\alpha\delta}{K_{14}} + \frac{\alpha\delta}{K_{23}} + \frac{\alpha\beta\delta}{K_{17}} + \frac{\alpha\beta\delta}{K_{26}} + \frac{\alpha\beta\gamma\delta}{K_{20}} + \frac{\alpha\beta\gamma\delta}{K_{29}}$$

$$\kappa_{PP2A} = \frac{\alpha}{K_3} + \frac{\alpha\beta}{K_6} + \frac{\alpha\beta\gamma}{K_9} + \frac{\delta}{K_{12}} + \frac{\alpha\delta}{K_{15}} + \frac{\alpha\delta}{K_{24}} + \frac{\alpha\beta\delta}{K_{18}} + \frac{\alpha\beta\delta}{K_{27}} + \frac{\alpha\beta\gamma\delta}{K_{21}} + \frac{\alpha\beta\gamma\delta}{K_{30}}$$

*Rate laws*

PKA

$$v_1 = \frac{k_1 \cdot PKA \cdot 0P}{K_1 \cdot \left(1 + \kappa_{PKA} - \frac{0P}{K_1}\right) + 0P} \qquad v_4 = \frac{k_4 \cdot PKA \cdot \alpha}{K_4 \cdot \left(1 + \kappa_{PKA} - \frac{\alpha}{K_4}\right) + \alpha} \qquad v_7 = \frac{k_7 \cdot PKA \cdot \alpha\beta}{K_7 \cdot \left(1 + \kappa_{PKA} - \frac{\alpha\beta}{K_7}\right) + \alpha\beta}$$

$$v_{22} = \frac{k_{22} \cdot PKA \cdot \delta}{K_{22} \cdot \left(1 + \kappa_{PKA} - \frac{\delta}{K_{22}}\right) + \delta} \qquad v_{25} = \frac{k_{25} \cdot PKA \cdot \alpha\delta}{K_{25} \cdot \left(1 + \kappa_{PKA} - \frac{\alpha\delta}{K_{25}}\right) + \alpha\delta} \qquad v_{28} = \frac{k_{28} \cdot PKA \cdot \alpha\beta\delta}{K_{28} \cdot \left(1 + \kappa_{PKA} - \frac{\alpha\beta\delta}{K_{28}}\right) + \alpha\beta\delta}$$

PKC

$$v_{10} = \frac{k_{10} \cdot PKC \cdot 0P}{K_{10} \cdot \left(1 + \kappa_{PKC} - \frac{0P}{K_{10}}\right) + 0P} \qquad v_{13} = \frac{k_{13} \cdot PKC \cdot \alpha}{K_{13} \cdot \left(1 + \kappa_{PKC} - \frac{\alpha}{K_{13}}\right) + \alpha} \qquad v_{16} = \frac{k_{16} \cdot PKC \cdot \alpha\beta}{K_{16} \cdot \left(1 + \kappa_{PKC} - \frac{\alpha\beta}{K_{16}}\right) + \alpha\beta}$$

$$v_{19} = \frac{k_{19} \cdot PKC \cdot \alpha\beta\gamma}{K_{19} \cdot \left(1 + \kappa_{PKC} - \frac{\alpha\beta\gamma}{K_{19}}\right) + \alpha\beta\gamma}$$

PP1

$$v_2 = \frac{k_2 \cdot PP1 \cdot \alpha}{K_2 \cdot \left(1 + \kappa_{PP1} - \frac{\alpha}{K_2}\right) + \alpha} \qquad v_5 = \frac{k_5 \cdot PP1 \cdot \alpha\beta}{K_5 \cdot \left(1 + \kappa_{PP1} - \frac{\alpha\beta}{K_5}\right) + \alpha\beta} \qquad v_8 = \frac{k_8 \cdot PP1 \cdot \alpha\beta\gamma}{K_8 \cdot \left(1 + \kappa_{PP1} - \frac{\alpha\beta\gamma}{K_8}\right) + \alpha\beta\gamma}$$

$$v_{11} = \frac{k_{11} \cdot PP1 \cdot \delta}{K_{11} \cdot \left(1 + \kappa_{PP1} - \frac{\delta}{K_{11}}\right) + \delta} \qquad v_{14} = \frac{k_{14} \cdot PP1 \cdot \alpha\delta}{K_{14} \cdot \left(1 + \kappa_{PP1} - \frac{\alpha\delta}{K_{14}}\right) + \alpha\delta} \qquad v_{17} = \frac{k_{17} \cdot PP1 \cdot \alpha\beta\delta}{K_{17} \cdot \left(1 + \kappa_{PP1} - \frac{\alpha\beta\delta}{K_{17}}\right) + \alpha\beta\delta}$$

$$v_{20} = \frac{k_{20} \cdot PP1 \cdot \alpha\beta\gamma\delta}{K_{20} \cdot \left(1 + \kappa_{PP1} - \frac{\alpha\beta\gamma\delta}{K_{20}}\right) + \alpha\beta\gamma\delta} \qquad v_{23} = \frac{k_{23} \cdot PP1 \cdot \alpha\delta}{K_{23} \cdot \left(1 + \kappa_{PP1} - \frac{\alpha\delta}{K_{23}}\right) + \alpha\delta} \qquad v_{26} = \frac{k_{26} \cdot PP1 \cdot \alpha\beta\delta}{K_{26} \cdot \left(1 + \kappa_{PP1} - \frac{\alpha\beta\delta}{K_{26}}\right) + \alpha\beta\delta}$$

$$v_{29} = \frac{k_{29} \cdot PP1 \cdot \alpha\beta\gamma\delta}{K_{29} \cdot \left(1 + \kappa_{PP1} - \frac{\alpha\beta\gamma\delta}{K_{29}}\right) + \alpha\beta\gamma\delta}$$

PP2A

$$v_3 = \frac{k_3 \cdot PP2A \cdot \alpha}{K_3 \cdot \left(1 + \kappa_{PP2A} - \frac{\alpha}{K_3}\right) + \alpha} \qquad v_6 = \frac{k_6 \cdot PP2A \cdot \alpha\beta}{K_6 \cdot \left(1 + \kappa_{PP2A} - \frac{\alpha\beta}{K_6}\right) + \alpha\beta} \qquad v_9 = \frac{k_9 \cdot PP2A \cdot \alpha\beta\gamma}{K_9 \cdot \left(1 + \kappa_{PP2A} - \frac{\alpha\beta\gamma}{K_0}\right) + \alpha\beta\gamma}$$

$$v_{12} = \frac{k_{12} \cdot PP2A \cdot \delta}{K_{12} \cdot \left(1 + \kappa_{PP2A} - \frac{\delta}{K_{12}}\right) + \delta} \qquad v_{15} = \frac{k_{15} \cdot PP2A \cdot \alpha\delta}{K_{15} \cdot \left(1 + \kappa_{PP2A} - \frac{\alpha\delta}{K_{15}}\right) + \alpha\delta} \qquad v_{18} = \frac{k_{18} \cdot PP2A \cdot \alpha\beta\delta}{K_{18} \cdot \left(1 + \kappa_{PP2A} - \frac{\alpha\beta\delta}{K_{18}}\right) + \alpha\beta\delta}$$

$$v_{21} = \frac{k_{21} \cdot PP2A \cdot \alpha\beta\gamma\delta}{K_{21} \cdot \left(1 + \kappa_{PP2A} - \frac{\alpha\beta\gamma\delta}{K_{21}}\right) + \alpha\beta\gamma\delta} \qquad v_{24} = \frac{k_{24} \cdot PP2A \cdot \alpha\delta}{K_{24} \cdot \left(1 + \kappa_{PP2A} - \frac{\alpha\delta}{K_{24}}\right) + \alpha\delta} \qquad v_{27} = \frac{k_{27} \cdot PP2A \cdot \alpha\beta\delta}{K_{27} \cdot \left(1 + \kappa_{PP2A} - \frac{\alpha\beta\delta}{K_{27}}\right) + \alpha\beta\delta}$$

$$v_{30} = \frac{k_{30} \cdot PP2A \cdot \alpha\beta\gamma\delta}{K_{30} \cdot \left(1 + \kappa_{PP2A} - \frac{\alpha\beta\gamma\delta}{K_{30}}\right) + \alpha\beta\gamma\delta}$$

*ODEs*

$$\frac{d}{dt}0P = v_2 + v_3 + v_{11} + v_{12} - v_1 - v_{10}$$

$$\frac{d}{dt}\alpha = v_1 + v_5 + v_6 + v_{14} + v_{15} - v_2 - v_3 - v_4 - v_{13}$$

$$\frac{d}{dt}\alpha\beta = v_4 + v_8 + v_9 + v_{17} + v_{18} - v_5 - v_6 - v_7 - v_{16}$$

$$\frac{d}{dt}\alpha\beta\gamma = v_7 + v_{20} + v_{21} - v_8 - v_9 - v_{19}$$

$$\frac{d}{dt}\delta = v_{10} + v_{23} + v_{24} - v_{11} - v_{12} - v_{22}$$

$$\frac{d}{dt}\alpha\delta = v_{13} + v_{22} + v_{26} + v_{27} - v_{14} - v_{15} - v_{23} - v_{24} - v_{25}$$

$$\frac{d}{dt}\alpha\beta\delta = v_{16} + v_{25} + v_{29} + v_{30} - v_{17} - v_{18} - v_{26} - v_{27} - v_{28}$$

$$\frac{d}{dt}\alpha\beta\gamma\delta = v_{19} + v_{28} - v_{20} - v_{21} - v_{29} - v_{30}$$

**Model 2** (Phenomenological model of increased α-dephosphorylation in presence of 2P/3P cMyBP-C)

All competition terms, rate laws and ODEs are identical to model 1, except for rate law $v_2$, which was multiplied with a 2P/3P-cMyB-C dependent activation term:

$v_2 = \left(\frac{k_2 \cdot PP1 \cdot \alpha}{K_2 \cdot \left(1 + \kappa_{PP1} - \frac{\alpha}{K_2}\right) + \alpha}\right) \cdot (1 + f_{act} \cdot h)$ , where the parameter $f_{act} \geq 0$ denotes a maximum activation factor and $h = \frac{r_{2P/3P}}{K_{act} + r_{2P/3P}}$ is a hyperbolic function of the relative amount of bis- and trisphosphorylated cMyBP-C $r_{2P/3P} = \frac{\alpha\beta + \alpha\delta + \alpha\beta\gamma + \alpha\beta\delta}{0P + \alpha + \alpha\beta + \alpha\beta\gamma + \delta + \alpha\delta + \alpha\beta\delta + \alpha\beta\gamma\delta}$, with a half-saturation constant $K_{act}$.

**Model 3** (Allosteric activation of α-dephosphorylation by 2P cMyBP-C)

All competition terms, rate laws and ODEs are identical to model 1, except for rate law $v_2$, which in model 3 accounts for allosteric activation by 2P:

$$v_2 = \frac{k_2 \cdot PP1 \cdot \alpha + k_A \cdot PP1 \cdot \alpha \cdot \frac{\alpha\beta + \alpha\delta}{\lambda K_A}}{K_2 + \frac{K_2 \cdot (\alpha\beta + \alpha\delta)}{K_A} + \frac{\alpha \cdot (\alpha\beta + \alpha\delta)}{\lambda K_A} + K_2 \cdot \left(\kappa_{PP1} - \frac{\alpha}{K_2}\right) + \alpha},$$

where $\lambda, k_A, K_A$ are parameters (see methods for details).

**Model 4**

*Substrate competition terms*

$$\kappa_{PKA} = \frac{0P}{K_1} + \frac{\alpha+\alpha'}{K_4} + \frac{\alpha\beta}{K_7} + \frac{\delta}{K_{22}} + \frac{\alpha\delta}{K_{25}} + \frac{\alpha\beta\delta}{K_{28}} \qquad \kappa_{PKC} = \frac{0P}{K_{10}} + \frac{\alpha+\alpha'}{K_{13}} + \frac{\alpha\beta}{K_{16}} + \frac{\alpha\beta\gamma}{K_{19}}$$

$$\kappa_{PP1} = \frac{\alpha}{K_2} + \frac{\alpha'}{K_{2,fast}} + \frac{\alpha\beta}{K_5} + \frac{\alpha\beta\gamma}{K_8} + \frac{\delta}{K_{11}} + \frac{\alpha\delta}{K_{14}} + \frac{\alpha\delta}{K_{23}} + \frac{\alpha\beta\delta}{K_{17}} + \frac{\alpha\beta\delta}{K_{26}} + \frac{\alpha\beta\gamma\delta}{K_{20}} + \frac{\alpha\beta\gamma\delta}{K_{29}}$$

$$\kappa_{PP2A} = \frac{\alpha}{K_3} + \frac{\alpha'}{K_{3,fast}} + \frac{\alpha\beta}{K_6} + \frac{\alpha\beta\gamma}{K_9} + \frac{\delta}{K_{12}} + \frac{\alpha\delta}{K_{15}} + \frac{\alpha\delta}{K_{24}} + \frac{\alpha\beta\delta}{K_{18}} + \frac{\alpha\beta\delta}{K_{27}} + \frac{\alpha\beta\gamma\delta}{K_{21}} + \frac{\alpha\beta\gamma\delta}{K_{30}}$$

*Rate laws*

PKA

$$v_1 = \frac{k_1 \cdot PKA \cdot 0P}{K_1 \cdot \left(1 + \kappa_{PKA} - \frac{0P}{K_1}\right) + 0P} \qquad v_4 = \frac{k_4 \cdot PKA \cdot \alpha}{K_4 \cdot \left(1 + \kappa_{PKA} - \frac{\alpha}{K_4}\right) + \alpha} \qquad v_{4,2} = \frac{k_{4,2} \cdot PKA \cdot \alpha'}{K_{4,2} \cdot \left(1 + \kappa_{PKA} - \frac{\alpha'}{K_{4,2}}\right) + \alpha'}$$

$$v_7 = \frac{k_7 \cdot PKA \cdot \delta}{K_7 \cdot \left(1 + \kappa_{PKA} - \frac{\delta}{K_7}\right) + \delta} \qquad v_{22} = \frac{k_{22} \cdot PKA \cdot \alpha\beta}{K_{22} \cdot \left(1 + \kappa_{PKA} - \frac{\alpha\beta}{K_{22}}\right) + \alpha\beta} \qquad v_{25} = \frac{k_{25} \cdot PKA \cdot \alpha\delta}{K_{25} \cdot \left(1 + \kappa_{PKA} - \frac{\alpha\delta}{K_{25}}\right) + \alpha\delta}$$

$$v_{28} = \frac{k_{28} \cdot PKA \cdot \alpha\beta\delta}{K_{28} \cdot \left(1 + \kappa_{PKA} - \frac{\alpha\beta\delta}{K_{28}}\right) + \alpha\beta\delta}$$

PKC

$$v_{10} = \frac{k_{10} \cdot PKC \cdot 0P}{K_{10} \cdot \left(1 + \kappa_{PKC} - \frac{0P}{K_{10}}\right) + 0P} \qquad v_{13} = \frac{k_{13} \cdot PKC \cdot \alpha}{K_{13} \cdot \left(1 + \kappa_{PKC} - \frac{\alpha}{K_{13}}\right) + \alpha} \qquad v_{13,2} = \frac{k_{13,2} \cdot PKC \cdot \alpha'}{K_{13,2} \cdot \left(1 + \kappa_{PKC} - \frac{\alpha'}{K_{13,2}}\right) + \alpha'}$$

$$v_{16} = \frac{k_{16} \cdot PKC \cdot \alpha\beta}{K_{16} \cdot \left(1 + \kappa_{PKC} - \frac{\alpha\beta}{K_{16}}\right) + \alpha\beta} \qquad v_{19} = \frac{k_{19} \cdot PKC \cdot \alpha\beta\gamma}{K_{19} \cdot \left(1 + \kappa_{PKC} - \frac{\alpha\beta\gamma}{K_{19}}\right) + \alpha\beta\gamma}$$

RSK2

$$v_{31} = \frac{k_{31} \cdot RSK2 \cdot 0P}{K_{31} + \frac{\delta}{K_{32}} + 0P} \qquad v_{32} = \frac{k_{32} \cdot RSK2 \cdot \delta}{K_{32} + \frac{0P}{K_{31}} + \delta}$$

PP1

$$v_2 = \frac{k_2 \cdot PP1 \cdot \alpha}{K_2 \cdot \left(1 + \kappa_{PP1} - \frac{\alpha}{K_2}\right) + \alpha} \qquad v_{2,fast} = \frac{k_{2,fast} \cdot PP1 \cdot \alpha'}{K_{2,fast} \cdot \left(1 + \kappa_{PP1} - \frac{\alpha'}{K_{2,fast}}\right) + \alpha'} \qquad v_5 = \frac{k_5 \cdot PP1 \cdot \alpha\beta}{K_5 \cdot \left(1 + \kappa_{PP1} - \frac{\alpha\beta}{K_5}\right) + \alpha\beta}$$

$$v_8 = \frac{k_8 \cdot PP1 \cdot \alpha\beta\gamma}{K_8 \cdot \left(1 + \kappa_{PP1} - \frac{\alpha\beta\gamma}{K_8}\right) + \alpha\beta\gamma} \qquad v_{11} = \frac{k_{11} \cdot PP1 \cdot \delta}{K_{11} \cdot \left(1 + \kappa_{PP1} - \frac{\delta}{K_{11}}\right) + \delta} \qquad v_{14} = \frac{k_{14} \cdot PP1 \cdot \alpha\delta}{K_{14} \cdot \left(1 + \kappa_{PP1} - \frac{\alpha\delta}{K_{14}}\right) + \alpha\delta}$$

$$v_{17} = \frac{k_{17} \cdot PP1 \cdot \alpha\beta\delta}{K_{17} \cdot \left(1 + \kappa_{PP1} - \frac{\alpha\beta\delta}{K_{17}}\right) + \alpha\beta\delta} \qquad v_{20} = \frac{k_{20} \cdot PP1 \cdot \alpha\beta\gamma\delta}{K_{20} \cdot \left(1 + \kappa_{PP1} - \frac{\alpha\beta\gamma\delta}{K_{20}}\right) + \alpha\beta\gamma\delta} \qquad v_{23} = \frac{k_{23} \cdot PP1 \cdot \alpha\delta}{K_{23} \cdot \left(1 + \kappa_{PP1} - \frac{\alpha\delta}{K_{23}}\right) + \alpha\delta}$$

$$v_{26} = \frac{k_{26} \cdot PP1 \cdot \alpha\beta\delta}{K_{26} \cdot \left(1 + \kappa_{PP1} - \frac{\alpha\beta\delta}{K_{26}}\right) + \alpha\beta\delta} \qquad v_{29} = \frac{k_{29} \cdot PP1 \cdot \alpha\beta\gamma\delta}{K_{29} \cdot \left(1 + \kappa_{PP1} - \frac{\alpha\beta\gamma\delta}{K_{29}}\right) + \alpha\beta\gamma\delta}$$

PP2A

$$v_3 = \frac{k_3 \cdot PP2A \cdot \alpha}{K_3 \cdot \left(1 + \kappa_{PP2A} - \frac{\alpha}{K_3}\right) + \alpha}$$

$$v_{3,fast} = \frac{k_{3,fast} \cdot PP2A \cdot \alpha'}{K_{3,fast} \cdot \left(1 + \kappa_{PP2A} - \frac{\alpha'}{K_{3,fast}}\right) + \alpha'}$$

$$v_6 = \frac{k_6 \cdot PP2A \cdot \alpha\beta}{K_6 \cdot \left(1 + \kappa_{PP2A} - \frac{\alpha\beta}{K_6}\right) + \alpha\beta}$$

$$v_9 = \frac{k_9 \cdot PP2A \cdot \alpha\beta\gamma}{K_9 \cdot \left(1 + \kappa_{PP2A} - \frac{\alpha\beta\gamma}{K_0}\right) + \alpha\beta\gamma}$$

$$v_{12} = \frac{k_{12} \cdot PP2A \cdot \delta}{K_{12} \cdot \left(1 + \kappa_{PP2A} - \frac{\delta}{K_{12}}\right) + \delta}$$

$$v_{15} = \frac{k_{15} \cdot PP2A \cdot \alpha\delta}{K_{15} \cdot \left(1 + \kappa_{PP2A} - \frac{\alpha\delta}{K_{15}}\right) + \alpha\delta}$$

$$v_{18} = \frac{k_{18} \cdot PP2A \cdot \alpha\beta\delta}{K_{18} \cdot \left(1 + \kappa_{PP2A} - \frac{\alpha\beta\delta}{K_{18}}\right) + \alpha\beta\delta}$$

$$v_{21} = \frac{k_{21} \cdot PP2A \cdot \alpha\beta\gamma\delta}{K_{21} \cdot \left(1 + \kappa_{PP2A} - \frac{\alpha\beta\gamma\delta}{K_{21}}\right) + \alpha\beta\gamma\delta}$$

$$v_{24} = \frac{k_{24} \cdot PP2A \cdot \alpha\delta}{K_{24} \cdot \left(1 + \kappa_{PP2A} - \frac{\alpha\delta}{K_{24}}\right) + \alpha\delta}$$

$$v_{27} = \frac{k_{27} \cdot PP2A \cdot \alpha\beta\delta}{K_{27} \cdot \left(1 + \kappa_{PP2A} - \frac{\alpha\beta\delta}{K_{27}}\right) + \alpha\beta\delta}$$

$$v_{30} = \frac{k_{30} \cdot PP2A \cdot \alpha\beta\gamma\delta}{K_{30} \cdot \left(1 + \kappa_{PP2A} - \frac{\alpha\beta\gamma\delta}{K_{30}}\right) + \alpha\beta\gamma\delta}$$

Isomerization

$$v_{iso,F} = k_{iso,F} \cdot \alpha' \qquad v_{iso,R} = k_{iso,R} \cdot \alpha$$

*ODEs*

$$\frac{d}{dt} 0P = v_2 + v_{2,fast} + v_3 + v_{3,fast} + v_{11} + v_{12} - v_1 - v_{10} - v_{31}$$

$$\frac{d}{dt} \alpha = v_1 + v_{31} - v_2 - v_3 - v_4 - v_{13} + v_{iso,F} - v_{iso,R}$$

$$\frac{d}{dt} \alpha' = v_5 + v_6 + v_{14} + v_{15} - v_{iso,F} + v_{iso,R} - v_{2,fast} - v_{3,fast} - v_{4,2} - v_{13,2}$$

$$\frac{d}{dt} \alpha\beta = v_4 + v_{4,2} + v_8 + v_9 + v_{17} + v_{18} - v_5 - v_6 - v_7 - v_{16}$$

$$\frac{d}{dt} \alpha\beta\gamma = v_7 + v_{20} + v_{21} - v_8 - v_9 - v_{19}$$

$$\frac{d}{dt} \delta = v_{10} + v_{23} + v_{24} - v_{11} - v_{12} - v_{22} - v_{32}$$

$$\frac{d}{dt} \alpha\delta = v_{13} + v_{13,2} + v_{22} + v_{26} + v_{27} + v_{32} - v_{14} - v_{15} - v_{23} - v_{24} - v_{25}$$

$$\frac{d}{dt} \alpha\beta\delta = v_{16} + v_{25} + v_{29} + v_{30} - v_{17} - v_{18} - v_{26} - v_{27} - v_{28}$$

$$\frac{d}{dt} \alpha\beta\gamma\delta = v_{19} + v_{28} - v_{20} - v_{21} - v_{29} - v_{30}$$
